# WUSCHEL modulates jasmonate signaling to control the balance between growth and defense in the shoot apical meristem

**DOI:** 10.64898/2026.02.10.705072

**Authors:** Pengfei Fan, Panagiotis Boumpas, Christian Wenzl, Yanfei Ma, Gernot Poschet, Jiao Zhao, Thomas Greb, Jan U. Lohmann

## Abstract

The growth-defense trade-off is an essential survival strategy for plants under environmental stress. However, how this balance is maintained at the meristematic level remains unclear. Here, we show that wounding of rosette leaves systemically inhibits inflorescence stem growth, associated with reduced cell proliferation in the inner cell layers of the shoot apical meristem (SAM), but not in the apical stem cell domain. Mechanistically, this selective cell behavior is dependent on the interplay between the growth-repressing jasmonate (JA) signaling and the growth-promoting stem cell regulator WUSCHEL (WUS). WUS mitigates wounding-induced growth inhibition by repressing JA-responsive gene expression via promoting the accumulation of JASMONATE-ZIM DOMAIN 3 (JAZ3), which encodes a JA signaling repressor. WUS on the one hand induces *JAZ3* mRNA accumulation and on the other hand directly competes with the JA receptor CORONATINE INSENSITIVE1 (COI1) for JAZ3 binding, thereby inhibiting JAZ3 protein degradation. Our findings identify a WUS-JAZ3-COI1 regulatory module that coordinates the growth-defense balance in the SAM upon systemic wounding, revealing a cell-type-specific mechanism for sustaining developmental robustness of the meristem during stress response.

## Introduction

To survive in a changing environment, plants must balance two essential tasks throughout their life cycles: sustaining intrinsic growth and defending against external threats. Typically, activation of defense responses occurs at the cost of growth, whereas prioritizing growth increases susceptibility to environmental stresses. The antagonistic relationship of growth and defense is widely recognized as the growth-defense trade-off^1^. Given that both disease resistance and crop yield are critical traits in plant breeding, understanding the mechanisms of this balance is essential for advancing agricultural productivity. While the growth-defense trade-off can be explained as a simple consequence of resource reallocation, emerging evidence suggests that the shift between growth and defense is actively regulated through crosstalk of multiple signaling pathways^2, 3^. Key components of growth regulatory modules, such as gibberellins (GA), brassinosteroids (BR), and light signaling pathways, have been shown to directly participate in the regulation of the growth-defense trade-off under stress conditions^2, 4–6^.

As the stem cell reservoir, the shoot apical meristem (SAM) plays an indispensable role in the growth regulation of all aboveground tissue in plants. The dynamic stem cell population in the SAM is maintained by a feedback loop involving the homeodomain transcription factor WUSCHEL (WUS) and the secreted peptide ligand CLAVATA3 (CLV3)^7–9^. *WUS* is expressed in the organizing center, while the synthesized protein migrates to the stem cells and activates *CLV3* expression^8, 10, 11^. In return, CLV3 represses *WUS* expression through signaling cascades mediated by the CLAVATA1, CLAVATA2 (CLV1, CLV2) and CLV2-CORYNE (CRN) complex^12, 13^. Despite a small surface-to-volume ratio compared to other plant organs, the SAM is capable of sensing environmental cues and integrating these signals into the internal growth regulatory network. Recent studies have shed light on the mechanisms of stem cell maintenance in the SAM in response to light, metabolic signals, reactive oxygen species (ROS), nitric oxide, heat stress, and viral infection, among others^14–20^. Notably, WUS-mediated inhibition of protein biosynthesis and salicylic acid (SA)-induced antiviral RNAi efficiently prevent virus replication in the SAM, suggesting that the SAM possesses specific strategies to counteract systemic infections^17, 19^.

In nature, plants frequently suffer from wounding stress mainly caused by herbivores. Upon wounding, first-wave signals, including electrical signals, calcium waves, and ROS, are propagated from the damaged tissue to the distal undamaged regions, hence inducing systemic wounding responses (SWR)^21^. Following this initial wave, the biosynthesis and signaling pathways of the phytohormone jasmonate (JA) are rapidly activated, thereby coordinating systemic defense and growth processes in response to SWR^22^. At the molecular level, JA signaling is initiated by the bioactive JA-isoleucine (JA-Ile)-dependent interaction of F-box protein CORONATINE INSENSITIVE1 (COI1) and the repressor proteins JASMONATE-ZIM DOMAINs (JAZs)^23–26^. This interaction leads to the degradation of JAZ proteins via the 26S proteasome, which releases MYC transcription factor proteins from JAZ-mediated repression^27–29^. Consequently, the expression of JA-responsive genes associated with defense and growth regulation is activated in response to wounding. Notably, JA signaling serves as a crucial regulatory node in reprogramming the growth-defense trade-off through the crosstalk with other pathways^6, 22, 30^. For example, the quintuple *JAZ* mutant *jazQ* displayed enhanced resistance to insect herbivores and fungal pathogens but exhibited inhibited growth and reduced fertility. However, mutation of *PHYB* in the *jazQ* mutant partially uncoupled this trade-off, resulting in the maintenance of robust growth of leaves, as well as effective defense^31^.

Wounding responses are globally induced, while the regulation of growth and development is highly tissue specific. How plants coordinate these processes to finetune the balance between growth and defense in a context-dependent manner remains largely unclear. Here, we used the SAM response to long-distance wounding signals as a model to investigate how the growth-defense trade-off is systemically coordinated in a tissue-specific manner. We show that long-term leaf wounding leads to a reduction of inflorescence stem growth and alters SAM morphology, which is mediated through the interplay between the WUS-CLV3 regulatory module and JA signaling pathways.

## Results

### Long-term leaf wounding inhibits inflorescence stem growth

To investigate the effects of systemic wounding on the shoot apex, we mechanically wounded rosette leaves with forceps after plants bolted (Fig. 1a). The treatment was repeated daily with the same degree of severity. Seven days after wounding treatment, wounded plants exhibited shorter inflorescence stems than unwounded plants (Fig. 1b, c), suggesting wounding leaves can systemically inhibit the growth of the inflorescence stem. As wounding treatment triggers the JA induction at the whole-plant level, we asked whether the wounding response includes activation of JA signaling in the shoot apex. To this end, we assessed the expression of four representative JA-responsive maker genes in the shoot apex as a readout of JA signaling activation in the wounding assay^29, 32, 33^. The expression of all markers was indeed increased three hours after wounding (Fig. 1d), indicating that wounding efficiently triggered the JA signaling in the shoot apex. To determine whether JA-related pathways are sufficient for inhibition of inflorescence stem growth observed after wounding, we applied JA directly on the shoot apex. Consistent with the phenotype observed after systemic wounding, JA-treated plants displayed shorter inflorescence stems compared to mock-treated plants (Fig. 1e, f). We then examined whether JA signaling is also required for wounding-induced stem growth inhibition by performing wounding treatment on *coi1-21* and *aos1-1* mutants, defective in JA signaling^34^ and JA biosynthesis^35^, respectively. To minimize the influence of flowering-time variation on inflorescence stem length measurement, all wounding treatments were initiated when the inflorescence stem was 0-0.5 cm in length. When assessed under these conditions, *coi1-21* and *aos1-1* plants displayed a significantly weaker reduction in inflorescence stem growth after wounding treatment compared to wild-type (WT) plants (Fig. 1g, h). Taken together, these results revealed that wounding of leaves triggers systemic defense responses and reduces inflorescence stem growth, mediated by the JA pathway. Thus, this assay provided us with a suitable model system to study the mechanisms of the growth-defense trade-off in the SAM.

**Fig. 1.**
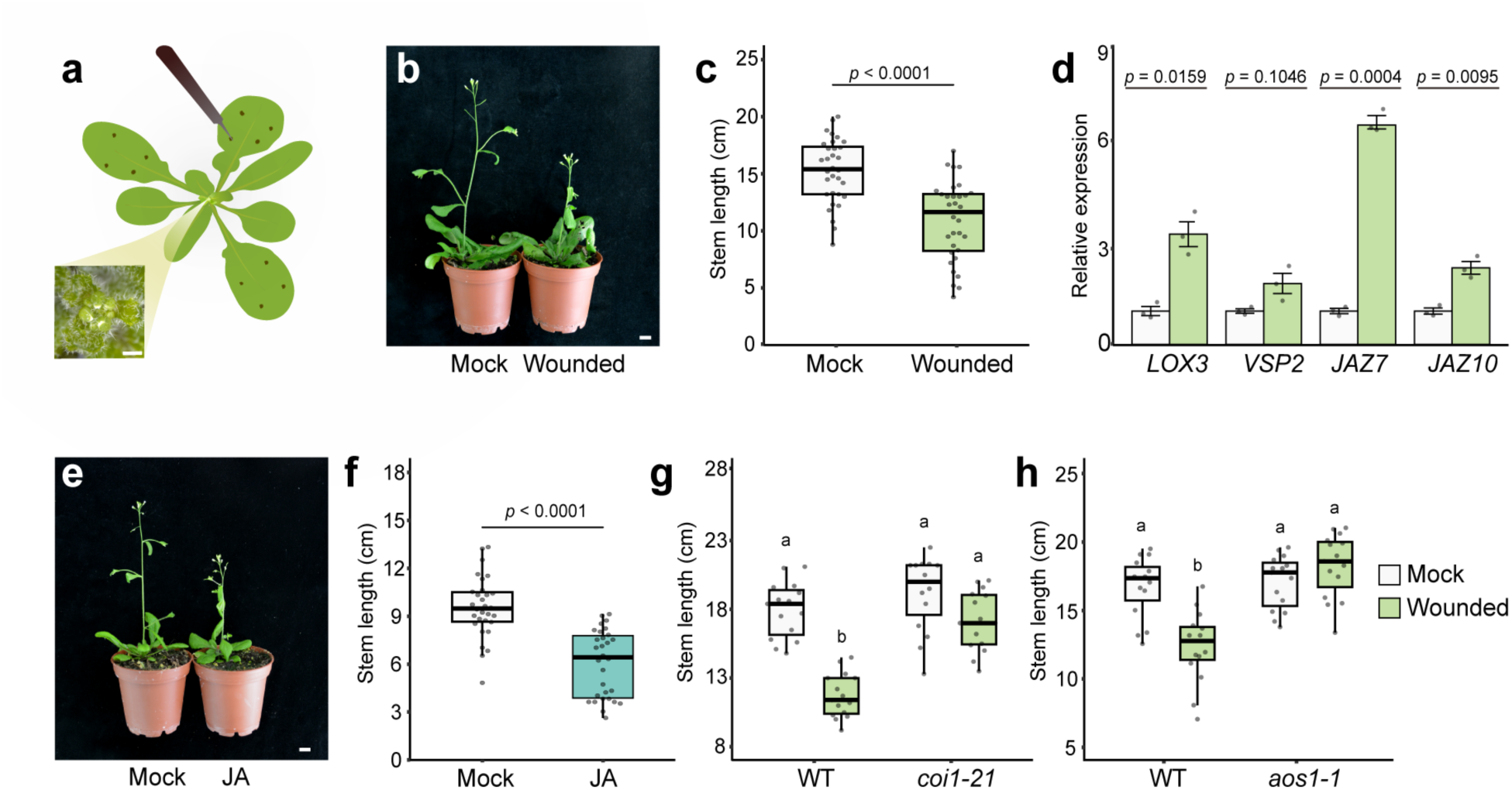
Systemic wounding and exogenous JA inhibit inflorescence stem growth. **a**, Wounding treatment in rosette leaves of *Arabidopsis*. When the main inflorescence stem was extended to 0-0.5 cm, wounding treatment was performed by punching four small holes per leaf with forceps. Three leaves were wounded each day. Scale bars, 1 mm. **b**, Representative plants after seven days of treatment. **c**, Quantification of inflorescence stem length after seven days of treatment. Data points are from three biological replicates (n ≥ 10 in each replicate per treatment). In all figures, “stem length” refers specifically to inflorescence stem length. For the boxplots in all figures, the central line indicates the median, the box represents the interquartile range of the data, and the whiskers extend to the lowest and highest values within the data. **d**, Expression of JA responsive marker genes in the SAM three hours after standard wounding treatment. Error bars represent the mean ± s.d from three biological replicates. **e**, Representative plants after five days of 50 μM JA treatment. **f**, Quantification of inflorescence stem length after five days of 50 μM JA treatment. Data points are from three biological replicates (n = 10 in each replicate per treatment). **g**, **h**, Quantification of inflorescence stem length of WT, *coi1-21* (**g**) and *aos1-1* (**h**) plants after seven days of wounding treatment. Statistical significance was analyzed using two-way ANOVA followed by Tukey’s multiple comparisons test (*p* < 0.05; n = 15 per condition). Three biological replicates were performed with similar results. Scale bars, 1cm (**b**, **e**). *P* values in **c**, **d**, and **f** were calculated using two-tailed Student’s t-tests.

### Systemic wounding and exogenous JA reshape SAM morphology

To explore how systemic wounding signals affect SAM homeostasis at the cellular level, we applied wounding or JA treatment to a triple reporter line (hereafter referred to as “*TR7*”) (Fig. 2a, b), which comprises three cell population markers, namely *pCLV3:mTagBFP2-NLS*, *pWUS:2xVENUS-NLS*, and *pUBQ10:3xmCherry-NLS*^36^. By combining 3D imaging and computational image analysis, we compared a series of morphological parameters of the SAM between wounded or JA-treated plants and mock-treated controls (Fig. 2c-g). SAMs were analyzed at identical inflorescence stem length to control for potential developmental delays. We found that both wounding and JA treatment caused a significant reduction in total cell number (Fig. 2j, k), while SAM size was only slightly decreased (Fig. 2h, i). Interestingly, the reduction of cell number occurred specifically in the inner cell layers, including part of peripheral zone and rib zone, whereas the cell number in the top two layers, where most of the stem cells reside, was not affected by wounding (Fig. 2l-o). These findings indicate that the inhibitory effects of wounding signals on the SAM are mainly restricted in the inner cell layers. In agreement with this observation, the number of *CLV3*-or *WUS*-expressing cells remained relatively constant after wounding (Fig. 2p, r), suggesting that stem cell maintenance is not disturbed by systemic wounding. However, exogenous JA treatment significantly reduced the numbers of *CLV3*- or *WUS*-expressing cells (Fig. 2q, s), which implies that the application of exogenous JA perturbs the stem cell regulatory system in contrast to systemic wounding. Moreover, the inhibitory effect of JA on SAM morphology is dose-dependent, as increasing the concentration of JA not only further reduced the SAM cell number and the number of *WUS-* or *CLV3*-expressing cells (Extended Data Fig. 1c, e, f, g), but also significantly diminished SAM size (Extended Data Fig. 1b). Notably, the cell number of the top two layers remained constant even under higher JA concentrations (Extended Data Fig. 1d), highlighting the robustness of this cell population to JA. Altogether, our findings indicate that systemic wounding signals modulate SAM morphology in the long-term by reducing the number of cells in the inner layers of the SAM without disturbing the stem cell regulatory system. Similarly, JA treatment preferentially targets the inner cell layers in a dose-dependent manner, but may further represses the WUS-CLV3 regulatory network at higher concentrations.

**Fig. 2.**
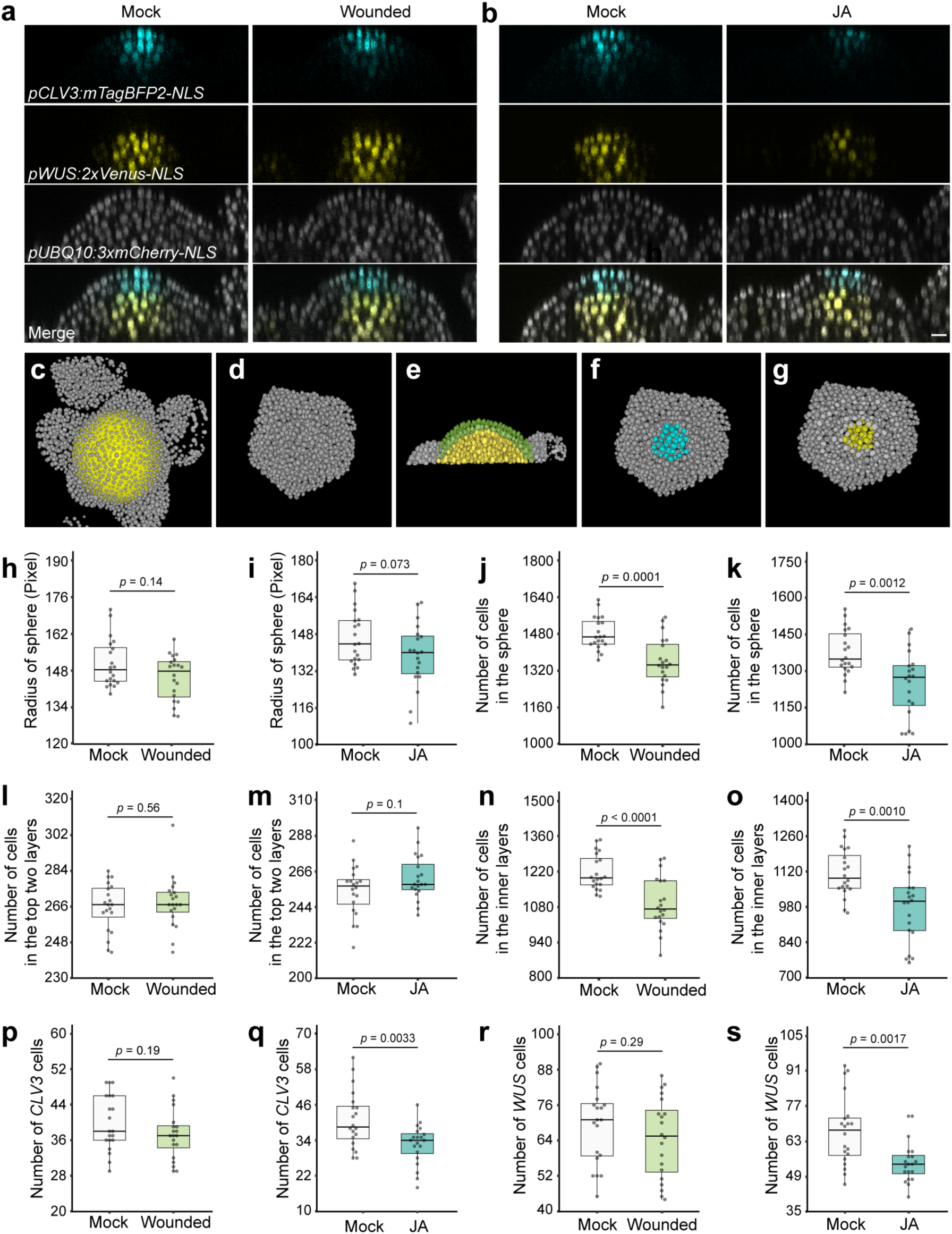
SAM morphology is modulated by systemic wounding and JA treatment. **a**,**b**, Representative central cross-sections of SAMs from the *TR7* triple reporter transgenic line after wounding (**a**) or treatment with JA (**b**). Scale bar, 10 μm. After systemic wounding, the SAMs were imaged when the inflorescence stem length was 10-12 cm. For JA treatment, SAMs were images after five days of treatment with 50 μM JA. **c**-**g**, Schematic representation of the SAM morphology-related parameters defined and assessed by the *TR7* computational image analysis pipeline: **c**, To determine the size of the SAM, a sphere (shown in yellow) was computationally fitted to the SAM; **d**, For the analysis of total SAM cell numbers, only cells in contact with the fitted sphere were retained to exclude floral organ primordia; **e**, The top two layers of the defined SAM area (shown in green) and the inner layers of the SAM beneath the top two layers (shown in yellow); **f**, domain of *CLV3*-expressing cells as identified by the *pCLV3:mTagBFP2-NLS* reporter (hereafter referred to as “*CLV3* cells”); **g**, domain of *WUS*-expressing cells as identified by the *pWUS:2xVenus-NLS* reporter (hereafter referred to as “*WUS* cells”). **h-i**, Quantification of SAM size after wounding (**h**) or JA treatment (**i**). SAM size is represented by the radius of the sphere defined in **c**. **j-k**, Quantification of the number of cells in the SAM after wounding (**j**) or JA treatment (**k**). The number of cells in the SAM is determined by counting the cells within the defined sphere. **l-m**, Quantification of the number of cells in the top two layers after wounding (**l**) or JA treatment (**m**). **n-o**, Quantification of the number of cells in the inner layers of the SAM after wounding (**n**) or JA treatment (**o**). Quantification of the number of *CLV3* cells or *WUS* cells s after wounding (**p**, **r**) or JA treatment (**q**, **s**). Data points are from two biological replicates (n = 10 per replicate). *P* values were calculated using two-tailed Student’s t-tests except **h**, in which Wilcoxon rank-sum test is used.

### WUS alleviates wounding-induced growth inhibition by repressing JA pathways

As systemic wounding stress did not significantly impair stem cell maintenance under our experimental conditions, we hypothesized that stem cells possess a certain degree of resilience to the effects of wounding stress, potentially through modulation of JA signaling by a SAM-specific regulatory machinery. Supporting our hypothesis, our previous RNA-seq analyses revealed that ubiquitous induction of WUS activity in seedlings results in differential expression of a large number of genes associated with JA signaling pathways^37^. To gain higher resolution at the tissue level, we profiled the transcriptome of dissected inflorescence SAMs from *pUBQ10:mCherry-GR-linker-WUS* plants following DEX-induced WUS activation. The newly generated RNA-seq dataset revealed distinct clustering of mock- and DEX-treated samples, as shown by PCA and Spearman correlation analysis (Extended Data Fig. 2a, b). A total of 3881 differentially expressed genes (DEG) were identified using a significant threshold of adjusted p-value < 0.05 and |log₂ fold change| > 0.5. Next, to delineate the distinct regulatory roles of WUS across different target gene groups, we performed Gene Ontology (GO) enrichment analysis separately on the upregulated and downregulated gene profiles. We found several JA-related GO terms, as well as the term “response to wounding”, significantly enriched among the gene with reduced expression (Fig. 3a), whereas no JA- or wounding-related GO terms were enriched among the genes with increased expression. Consistently, most identified differentially expressed genes associated with JA pathways were reduced upon induction of ubiquitous WUS activity (Extended Data Fig. 2c). Altogether, the transcriptome profiles underline that WUS may function as a negative regulator of JA response within the SAM.

**Fig. 3.**
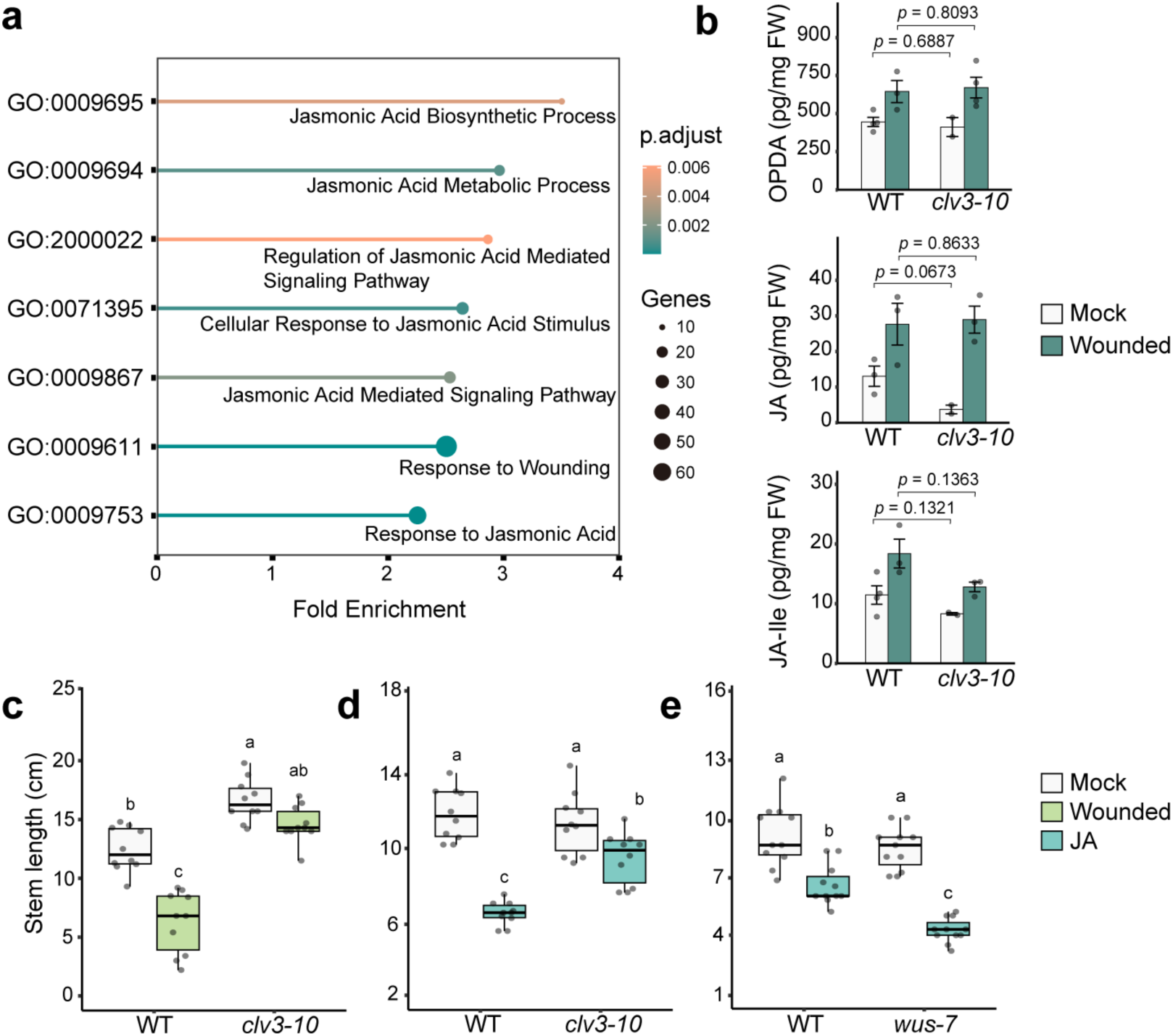
WUS regulates JA pathway in the SAM. **a**, Enrichment of JA- and wounding-related GO terms among negatively regulated WUS target genes identified by RNA-seq in SAMs with ubiquitous induction of WUS activity. SAMs of *pUBQ10:mCherry-GR-WUS* plants were collected for RNA-seq analysis two hours after mock or 10 μM DEX treatment. GO analysis was performed on genes with adjusted p-value (p._adj_) < 0.05 and |Log_2_FC| > 0.5. Three biological replicates were analyzed. All the significantly enriched GO terms are presented in the supplementary documents. **b**, Quantification of OPDA, JA, and JA-Ile contents in the shoot apexes of WT and *clv3-10* plants three hours after wounding treatment. Error bars indicate the mean ± s.d. of results from two to four biological replicates. *P* values were calculated using Welch’s t-test. **c**, Quantification of inflorescence stem length of WT and *clv3-10* plants after seven days of wounding treatment (n = 10 per condition). **d**, Quantification of inflorescence stem length of WT and *clv3-10* plants after seven days of mock or 50 μM JA treatment (n = 10 per condition). **e**, Quantification of inflorescence stem length of WT *and wus-7* plants after seven days of mock or 50 μM JA treatment (n = 11 per condition). Statistical significance in **c**-**e** was analyzed using two-way ANOVA followed by Tukey’s multiple comparisons test (*p* < 0.05). Data points presented in **c**-**e** are from one biological replicate. Three biological replicates were performed for **c** and **d,** and two for **e**, all with similar results.

In the JA biosynthesis pathway, OPDA (12-oxo-phytodienoic acid) serves as a precursor of JA, which is subsequently converted into the bioactive hormone JA-Ile^28^. Given that WUS potentially represses the expression of genes involved in JA synthesis, we asked whether enhancing WUS activity in the shoot apex could attenuate JA accumulation induced by systemic wounding. To test this, we quantified concentrations of OPDA, JA and JA-Ile in the shoot apices of WT plants and the *clv3-10* mutants, in which loss of CLV3 function results in elevated WUS activity^38^. Under unwounded conditions, the levels of JA and JA-Ile were lower in the shoot apex of *clv3-10* mutants compared to WT plants, while OPDA concentrations remained similar (Fig. 3b). However, the concentration of all the three metabolites increased in both WT and *clv3-10* upon wounding with a slight attenuation observed for JA-Ile (Fig. 3b). These results suggest that elevated WUS activity suppresses basal JA accumulation, whereas the JA biosynthesis pathway remains responsive to systemic wounding, even in the *clv3-10* mutant background.

Next, to test whether WUS mitigates the stem growth inhibition caused by wounding signals, we conducted a seven-day JA or wounding treatment in both *clv3-10* and *wus-7* mutants, the latter of which represents a weak loss-of-function allele of *WUS*. Compared to WT plants, the inflorescence stem growth of the *clv3-10* mutants was more resistant to systemic wounding treatment and exogenous JA (Fig. 3c, d), whereas *wus-7* mutants were more sensitive and displayed a higher level of inflorescence growth inhibition upon exogenous JA (Fig. 3e). Altogether, these results reveal a dual role of WUS in the regulation of wounding response: repressing JA signaling and biosynthesis in the SAM at steady state, and alleviating inflorescence stem growth inhibition upon wounding.

### WUS promotes *JAZ3* expression in the SAM

JA signaling regulation is tissue specific and relies on distinct signaling cassettes in different tissues^32, 39^. Accordingly, to explore the molecular basis of how WUS modulates the wounding response, we first characterized the expression pattern of canonical JA signaling components in the shoot apex. 3D imaging of *pCOI1:COI1-VENUS* reporter line^40^ showed that the JA receptor COI1 accumulated ubiquitously in the SAM (Extended Data Fig. 3a). To examine the expression of *MYC* genes, which encode key transcription factors regulating the JA response, we performed in situ hybridization to visualize RNA accumulation patterns of *MYC2/3/4/5*. Under our conditions, only *MYC3* expression was detectable in the SAM (Extended Data Fig. 3b), with enrichment in the rib zone. The low or undetectable expression for other *MYC* genes may be due to technical limitations, such as probe efficiency. Nevertheless, these results indicate that at least *MYC3* is expressed in the SAM.

Given that WUS globally suppresses JA signaling, we next examined whether *JAZ* genes, encoding the core repressors of JA signaling pathway, were involved in WUS-mediated regulation in response to wounding signals. Since our SAM transcriptome profiling identified only *JAZ6* as differentially expressed gene upon induction of ubiquitous WUS activity, we revisited our RNA-seq dataset from seedlings to gain a more comprehensive view of potential WUS targets^37^ (Extended Data Fig. 3c). Among all the *JAZ* genes potentially regulated by WUS, we checked the expression pattern of *JAZ2, JAZ3, JAZ5* and *JAZ6* by in situ hybridization. In our settings, only *JAZ3* and *JAZ5* mRNAs were detectable in the SAM, whereas no signal was observed with *JAZ2 or JAZ6* probes (Extended Data Fig. 3d). As part of an intricate feedback regulation, *JAZ* expression can be activated by JA treatment^33^. We therefore tested whether *JAZ* expression in the SAM was affected by JA treatment. Indeed, the expression of *JAZ2*, *JAZ3*, and *JAZ5* was significantly enhanced two hours after JA application (Extended Data Fig. 3d).

As *JAZ3* expression was readily detectable under basal conditions, we focused on this key regulator to explore the molecular basis of the interaction between WUS and JA signaling. Notably, WUS was shown to bind the *JAZ3* promoter and induction of ubiquitous WUS activity in seedlings led to increased *JAZ3* expression ^20, 37^(Fig. 4a, b). In line with this, *JAZ3* expression was markedly increased in the shoot apex of *clv3-10* mutants compared to WT plants (Fig. 4c). Moreover, eChIP-qPCR revealed significant WUS enrichment at four of the five examined sites within the *JAZ3* promoter, with the highest enrichment observed near the transcription start site and within the 5′-UTR, suggesting that WUS binds to the *JAZ3* locus to directly regulate its expression (Extended Data Fig. 4). In addition, in situ hybridization and 3D imaging of the *pJAZ3:GFP-NLS* reporter showed that *JAZ3* is specifically expressed in the center of the SAM and young primordia, with enriched signal in the organizing center and central rib zone (Figure 4d, e). Furthermore, consistent with previous results, increased RNA accumulation was observed in the SAM two hours after ubiquitous induction of WUS activity (Fig. 4f). Collectively, these findings demonstrate that WUS binds the *JAZ3* locus and directly activates its expression in the SAM. The finding that WUS induces the expression of a key negative regulator of JA signaling fits well with the observation that overall, WUS leads to a reduction in expression of JA responsive genes.

**Fig 4:**
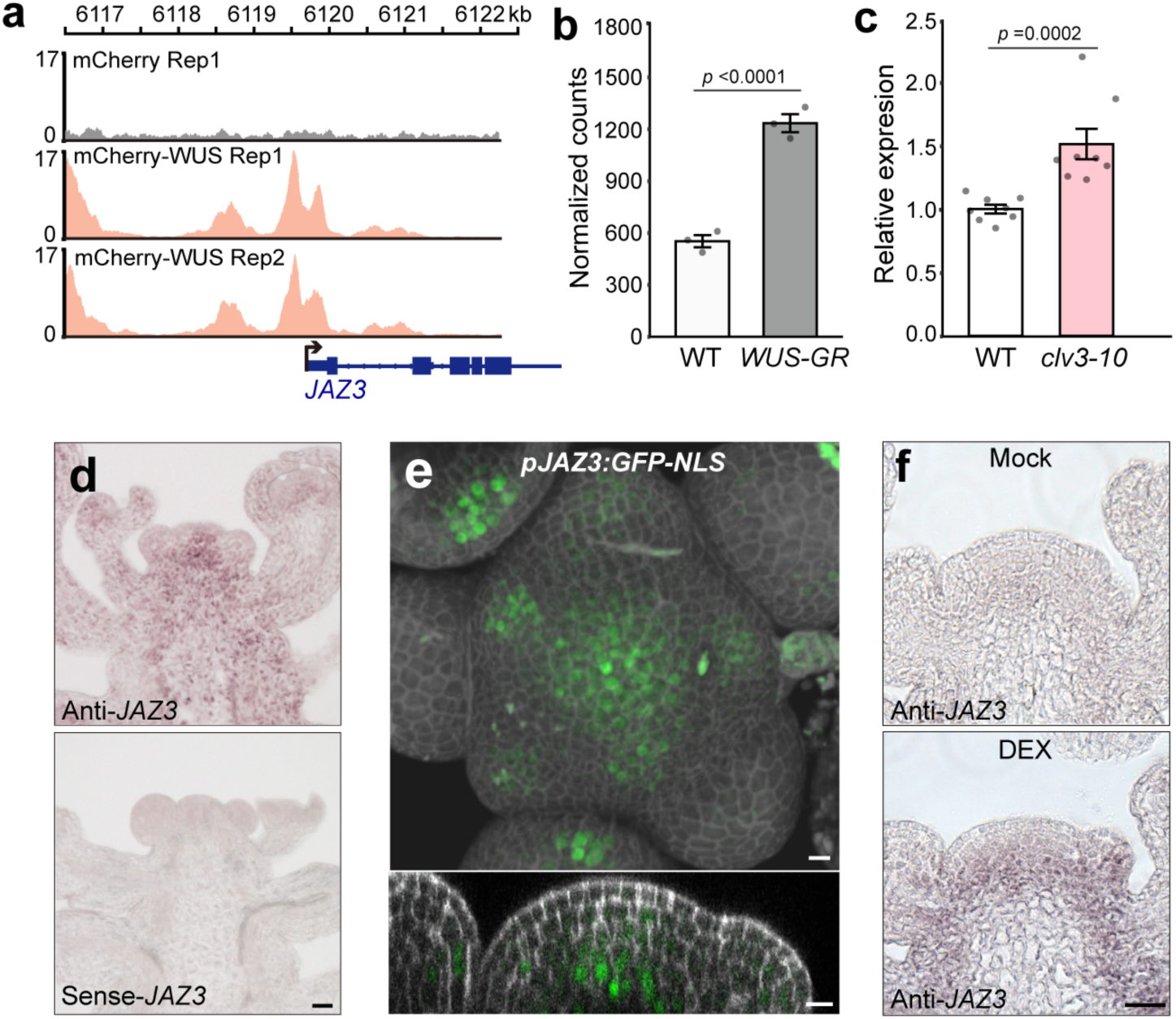
WUS directly promotes *JAZ3* expression. **a**, Visualization of eChIP-seq reads along the promoter regions of *JAZ3* identified in DEX treated *pUBQ10:mCherry-GR* and *pUBQ10:mCherry-GR-linker-WUS* seedlings. Bottom: schematic of the partial *JAZ3* (AT3G17860.1) genomic region. Black arrow marks the transcription start site and direction. Blue lines indicate introns. Thicker boxes indicate exons and the thinner box represents the 5 ’untranslated region. **b**, Normalized counts of *JAZ3* mRNA detected in RNA-seq data of DEX-treated WT (Col-0) and *pUBQ10:mCherry-GR-linker-WUS* (*WUS-GR*) *seedlings.* Error bars indicate the mean ± s.d. of results from three biological replicates. **c**, mRNA levels of *JAZ3* in shoot apexes of WT or *clv3-10* plants as measured by RT-qPCR. Error bars represent the mean ± s.d. from eight biological replicates. *P* values were calculated using Wilcoxon rank-sum test. **d**, *JAZ3* RNA accumulation pattern in the SAM as detected by in situ hybridization. Scale bar, 25 μm. **e**, Representative 3D volume view and the central cross-section of the SAM in *pJAZ3:GFP-NLS* plants. The gray channel shows cell walls stained by 4’,6-diamidino-2-phenylindole (DAPI). Scale bar, 10 μm. SAMs were imaged when the inflorescence stem length was 5-8 cm. **f**, *JAZ3* RNA accumulation in the SAM of *pUBQ10:mcherry-GR-linker-WUS* plants after two hours of mock or DEX treatment, detected by in situ hybridization. Shoot apexes of *pUBQ10:mcherry-GR-linker-WUS* plants were sprayed with 10 μM DEX on the shoot apex. SAMs were treated and collected when the stem length of the plants was 5-8 cm. Scale bar, 25 μm. The signal detection reaction was stopped earlier than in **d** to avoid signal oversaturation (n = 4). Experiments were independently performed two times with similar results.

### WUS competes with COI1 for interaction with JAZ3

Given that JAZ3 acts as a transcriptional repressor and its expression pattern largely overlaps with the WUS expression domain in the SAM, we asked whether in addition to transcriptional regulation, WUS may modulate JA signaling through physically interacting with JAZ3. Indeed, we found the two proteins to interact in a yeast-two hybrid assay (Y2H) (Fig. 5a). Since WUS contains intrinsically disordered domains that tend to cause unspecific interactions, we sought to validate this finding *in planta* by two complementing approaches. Using transient co-expression in *Nicotiana benthamiana* leaves we first applied TurboID-based proximity labeling. JAZ3 fused to miniTurbo was able to biotinylate WUS-GFP, but not GFP-NLS as detected following immunoprecipitation with GFP antibody (Fig. 5b), demonstrating that JAZ3 and WUS come in close contact in living plant cells. Next, we used Acceptor Photobleaching Förster Resonance Energy Transfer (AP-FRET) to identify interacting domains in both proteins. AP-FRET not only confirmed the WUS-JAZ3 interaction, but also demonstrated that both the ZIM domain and the C-terminal Jas domain in JAZ3 are required for its interaction with WUS (Fig. 5c, d). The Jas motif in JAZ proteins is essential for COI1-dependent 26S proteasome degradation^23, 25^. Interestingly, it has been reported that two conserved positively charged amino acids within the Jas domain are required for JAZ-COI interaction^41^ and mutating these two residues in JAZ3 abolished its interaction with WUS (Extended Data Fig. 5). For the WUS protein, the unstructured region within its dimerization domain was required for the interaction with JAZ3, as deletion of this domain abolished the interaction (Fig. 5f).

**Fig. 5.**
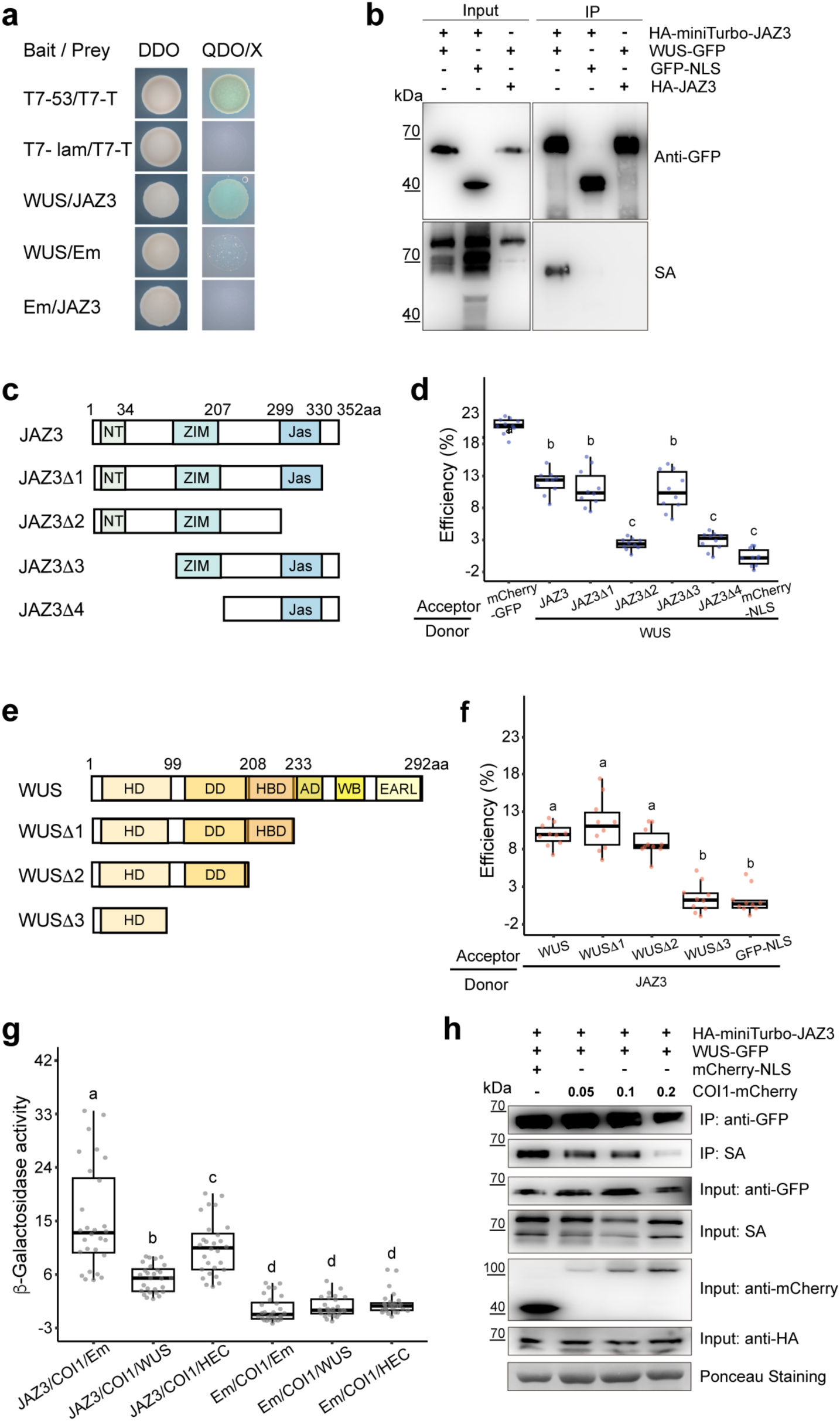
WUS competes with COI1 for binding to JAZ3. **a**, Yeast two-hybrid assay showing the interaction between WUS and JAZ3. DDO (double dropout medium): SD/-Trp/-Leu. QDO/X (quadruple dropout medium supplemented with x-α-gal): SD/-Trp/-Leu/-His/-Ade/x-α-gal. Bait: proteins fused to binding domain (BD). Prey: proteins fused to activation domain (AD). T7-53 /T7-T was used as positive control, and T7-lam/ T7-T as negative control. Em: Empty vector. Experiments were independently repeated three times with similar results. **b**, TurboID-based proximity labeling assay showing the interaction between WUS and JAZ3 upon transient co-expression in *N. benthamiana* leaves. Biotin (50 μM) was infiltrated into leaves 30 minutes before sample collection at 2 days post-agroinfiltration (2 dpa). Total proteins were extracted and immunoprecipitated using GFP-Trap magnetic beads. Input and IP fractions were analyzed by immunoblotting with anti-GFP and Streptavidin (SA). Three biological replicates were performed with similar results. **c**, **e**, Schematic representation of truncated versions of JAZ3 (**c**) or WUS (**e**). Numbers indicate the amino acid (aa) positions where truncations begin or end. HD: Homeodomain; DD: Dimerization domain; HBD: HAM binding domain; AD: Acidic domain; WB: WUS-box; EARL: EAR-like domain. **d**, **f**, AP-FRET experiments identifying the domains in WUS and JAZ3 required for their interaction upon transient co-expression in *N. benthamiana* leaves. (**d**) Interaction of WUS with truncated variants of the JAZ3 protein; (**f**) Interaction of JAZ3 with truncated variants of the WUS protein. GFP was C-terminally fused to WUS (and truncated variants), and mCherry was N-terminally fused to JAZ3 (and truncated variants). A mCherry-GFP fusion was used as positive control. Data points represent the FRET efficiencies measured from nuclei (n = 10). Statistical significances in **d** and **f** were analyzed using one-way ANOVA followed by Tukey’s multiple comparisons test (*p* < 0.05). Three biological replicates were performed with similar results. **g**, Yeast three-hybrid (Y3H) assay showing the interaction between JAZ3 and COI1 in the presence or absence of WUS. JAZ3 fused to the AD was used as prey, and COI1 fused to the BD was used as bait. β-galactosidase activity was measured in co-transformed yeast cells grown in liquid triple dropout medium (SD/-Trp/-Leu/-Met) supplemented with 40 μM coronatine for three days. Em, empty vector. The HECATE1 (HEC) bHLH transcription factor was used as negative control. Each data point represents an independent co-transformed yeast colony (n = 28). Statistical significance was analyzed using one-way ANOVA followed by Tukey’s multiple comparisons test (*p* < 0.05). **h**, JAZ3-WUS interaction under varying COI1 levels, as shown by TurboID-based proximity labeling assay. Constructs expressing HA-miniTurbo-JAZ3, WUS-GFP, and either COI1-mCherry or mCherry-NLS were transiently co-transformed into *N. benthamiana* leaves. The level of COI1-mCherry expression was modulated by adjusting the OD_600_ of infiltrated *agrobacterium* cultures. Total proteins were extracted and immunoprecipitated using GFP-Trap magnetic beads. Input and IP fractions were analyzed by immunoblotting with anti-GFP, SA, anti-mCherry, and anti-HA. Three biological replicates were performed with similar results.

Since WUS and COI1 share binding sites in the JAZ3 protein, we wondered whether WUS may compete with COI1 for binding to JAZ3. Such interaction would protect the JAZ3 protein from degradation in response to JA signaling, which would lead to a reduction in JA response. Indeed, we found that the interaction between JAZ3 and COI1 was significantly reduced in the presence of WUS in a yeast-three hybrid assay (Fig. 5g). However, this reduction was rescued by higher levels of JA-Ile (Extended Data Fig. 6), suggesting that JA influences the competition between WUS and COI1 for JAZ3 binding. Similarly, TurboID-based proximity labeling assays confirmed that COI1 interfered with the WUS-JAZ3 interaction in a dose-dependent manner (Fig. 5h). Collectively, these results suggest that the interplay between growth and defense regulation in response to wounding signals is potentially mediated by the regulation of JAZ3 protein accumulation by WUS through two converging mechanisms, namely transcriptional activation and protection from COI1 mediated degradation.

### JAZ3 regulates inflorescence stem growth in response to systemic wounding

Given that JAZ3 functions as a key regulatory node of growth-defense balance in the shoot apex upon wounding stress, we next asked whether it also participates in regulating the inflorescence stem growth trade-off under wounding conditions. To test this, we generated *pWUS:JAZ3* transgenic lines under Col-0 and *TR7* background, in which *JAZ3* expression is enhanced in the *WUS*-expressing domain. *pWUS:JAZ3* plants exhibited an early flowering phenotype compared to control plants, characterized by a slightly earlier bolting time and a reduced number of rosette leaves (Extended Data Fig. 7a, b). Next, to evaluate the effects of *JAZ3* overexpression on SAM morphology, we performed 3D imaging and computational analysis of SAMs from *pWUS:JAZ3*/*TR7* plants. The SAM of *pWUS:JAZ3* displayed an overall slightly increased cell number and SAM size but similar number of *WUS*- or *CLV3*-expressing cells as *TR7* (Extended Data Fig. 7c-h). Moreover, *JAZ3* expression under the *WUS* promoter failed to rescue the fasciated inflorescence shoot apex phenotype of the *clv3-10* mutant (Extended Data Fig. 7i). Altogether, these results suggest that overexpressing *JAZ3* does not significantly affect stem cell maintenance in the SAM but promotes floral transition. To further decipher the function of JAZ3 in the SAM, we generated *jaz3-cr* mutant using the *CRISPR/Cas9* knockout system^42^ (Extended Data Fig. 8a). *Jaz3-cr* plants exhibited WT-like growth and developmental phenotypes, but delayed bolting time similar as the *jaz3/5/10* triple mutant^43^ (Extended Data Fig. 8b, c).

Next, we performed systemic wounding treatments on *pWUS:JAZ3* and *jaz3-cr* plants. *pWUS:JAZ3* transgenic lines exhibited reduced stem growth inhibition compared with WT plants upon wounding, although the degree of this reduced sensitivity varied among different transgenic lines (Fig. 6a), indicating that elevating *JAZ3* expression can partially reduce the growth trade-off under wounding conditions. By contrast, stem growth of *jaz3-cr* was more sensitive to systemic wounding (Fig. 6b). Taken together, these results show that JAZ3, like WUS, acts as a positive regulator in alleviating JA-mediated inhibition of inflorescence stem growth under systemic wounding.

**Figure 6.**
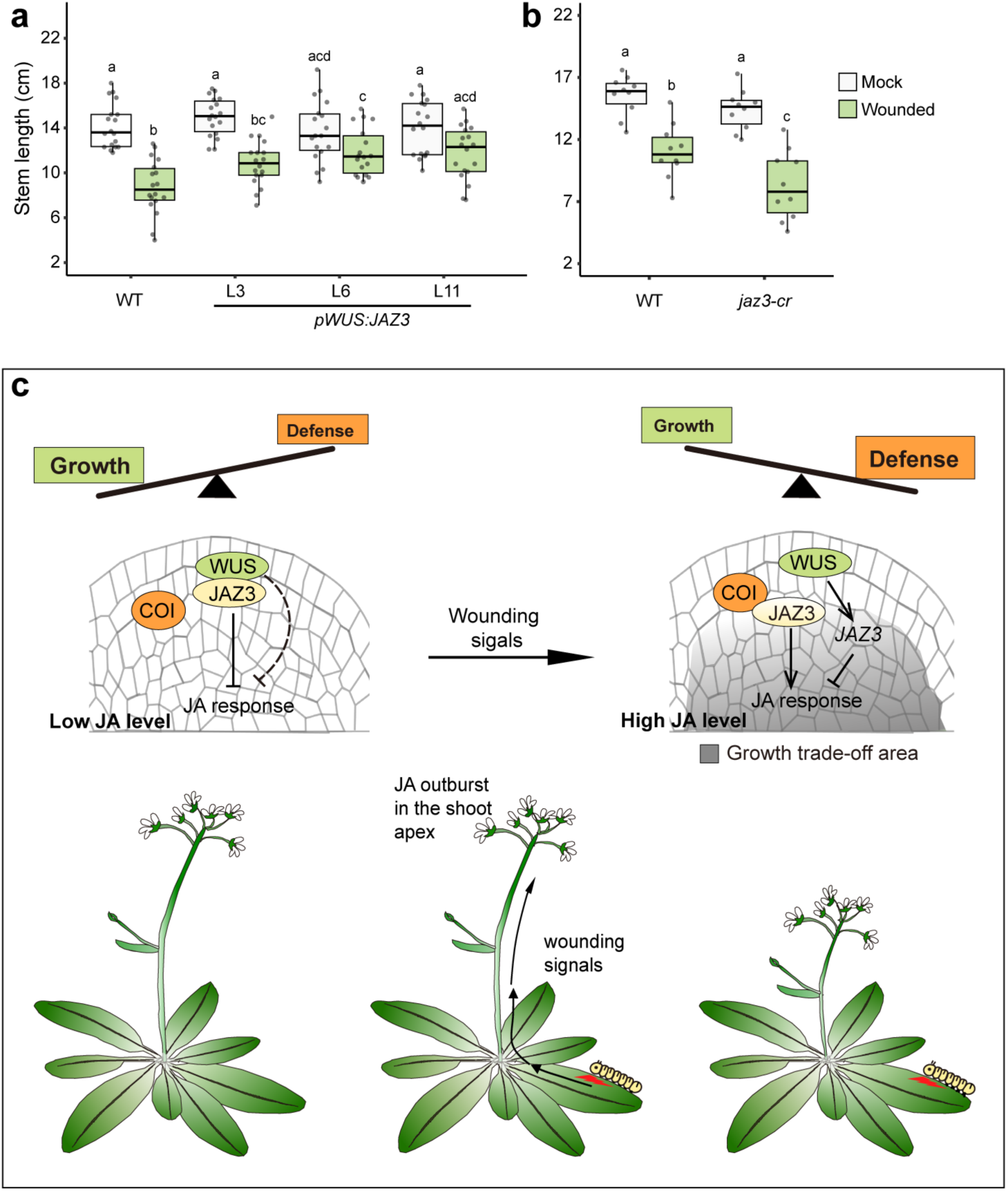
JAZ3 regulates inflorescence stem growth in response to JA-mediated systemic wounding signals. **a**, **b**, Quantification of inflorescence stem length of WT, *pWUS:JAZ3* (**a**), and *jaz3-cr* (**b**) after seven days of wounding treatment (n = 10-18 per condition). Statistical significance was analyzed using two-way ANOVA followed by Tukey’s multiple comparisons test (*p* < 0.05). Two independent biological replicates were performed with similar results. **c**, A proposed model depicting the growth-defense trade-off regulation in the SAM in response to systemic wounding signals. Under steady-state conditions, where growth is prioritized over defense, low JA levels in the SAM allow JAZ3 to preferentially interact with WUS rather than COI1. This WUS-JAZ3 interaction suppresses JA signaling, thereby maintaining stem cell homeostasis in the SAM (left panel). Upon leaf wounding, systemic wounding signals reach the shoot apex and trigger a JA burst in the SAM. Elevated JA levels enable COI1 to outcompete WUS for interacting JAZ3, leading to JAZ3 degradation (right panel). Meanwhile, WUS promotes *JAZ3* expression and represses JA-responsive gene expression, which in turn confers resilience to wounding stress in the stem cell region. As a result, cell proliferation in the inner layers of the SAM and inflorescence stem growth are inhibited in response to systemic wounding signals.

## Discussion

Since wounding responses are coordinated across the entire plant, regulation of the growth-defense balance is expected at the whole-plant level. However, how this trade-off is manifested in unwounded distal tissues remains largely unknown. Here, we propose a tissue-specific model for the growth-defense trade-off in the SAM upon systemic wounding. In this model, the systemic wounding signals originating from injured rosette leaves reach the SAM and remodel its morphology. The cell population in the top two layers are largely protected from wounding effects, thereby preserving the overall reproductive potential of the plant. In contrast, a reduction in cell number within the inner cell layers leads to a growth penalty in the inflorescence stem. This tissue-specific growth trade-off pattern is governed by the interplay between JA signaling and a novel WUS-mediated regulatory module. WUS promotes *JAZ3* expression, thereby reinforcing repression of JA signaling within the stem cell niche. In addition, WUS competes with the JA receptor COI1 for JAZ3 binding, protecting the latter from degradation. Together, the WUS-JAZ3-COI1 module functions as a molecular switch that toggles the SAM between growth and defense states upon wounding.

Wounding in the leaves of *Arabidopsis* seedlings has been reported to inhibit the growth of unwounded root^44^. In our study, we found that wounding rosette leaves retards inflorescence stem growth and reduces cell numbers in the SAM, further confirming the profound impact of systemic wounding on overall plant growth and development. At the cellular level, a key question underlying this growth trade-off is whether it results primarily from the inhibition of cell division, cell elongation, or both. Previous studies have shown that leaf growth inhibition upon wounding is mainly due to reduced cell division^39^. On the other hand, exogenous JA application suppresses primary root growth by repressing both processes^45^. In the context of stem development, elongation is primarily driven by cell proliferation in the SAM rib zone and, in the internode, it proceeds sequentially through slow cell division followed by rapid expansion^46^. Our results suggest that the inhibition of cell division in the basal cell layers of the SAM underlies the reduced stem growth observed upon systemic wounding. However, a potential contribution of altered cell elongation in the internode cannot be excluded, as the change of the cell size was not evaluated in our study. These findings indicate that the growth trade-off induced by systemic wounding displays cell-type specificity and hence is regulated in a highly context-dependent manner.

In the inflorescence SAM, the top two cell layers contain stem cells and progenitor cells of the epidermis and lateral organs^47^. As the cell number of the top two layers of the SAM remains unchanged upon wounding, this raises the possibility that these layers are either insulated from the inhibitory effects of wounding or that such effects are rapidly neutralized upon their entry. Given that *JAZ* genes are well-established downstream markers of the JA pathway^33^, their expression pattern serves as a useful readout of JA signaling in plant tissues. In our study, the JA-induced expansion of *JAZ2*, *JAZ3*, and J*AZ5* expression into the top two layers supports the latter hypothesis, suggesting that systemic signals do reach these layers but are usually attenuated. Moreover, systemic wounding did not significantly alter the number of *WUS*- or *CLV3*-expressing cells. By contrast, exogenous JA application led to a clear reduction in both types of cells, indicating the WUS-CLV3 regulatory system can be repressed by exogenous JA. Notably, this negative effect is dose-dependent, as increasing concentrations of exogenous JA aggravated the reduction. Considering that the JA burst induced by systemic wounding may be less intense than an exogenous JA application, the milder impact of systemic wounding on *WUS*- and *CLV3*-expressing cell numbers suggests that SAM activity can recover from rhythmic, wounding-associated JA stress. However, when JA levels exceed a certain threshold in levels or duration, stem cell activity is affected. Collectively, these results highlighted the robustness of the stem cell regulatory system in buffering against wounding stress and pointed to a stem cell specific mechanism that counteracts the inhibitory effects of both systemic wounding and JA signaling.

Consistent with this idea, our study further identified a molecular mechanism in which WUS functions as a key component of the stem cell resilience to wounding by modulating JA responses. Transcriptomic profiling shows that WUS globally represses the expression of genes involved in JA biosynthesis, signaling and response. Consistent with the reduction of JA biosynthesis genes by WUS, we found JA levels to be reduced in the shoot apex of *clv3-10* mutant plants, in which *WUS* expression is massively expanded. Although the levels of bioactive JA-Ile were only slightly reduced in the shoot apex of *clv3-10* mutants following leaf wounding, the mutant exhibited significantly greater long-term resistance to stem growth inhibition. One possible explanation is that enhanced WUS activity in the *clv3-10* mutant more efficiently terminates JA signaling in the stem cell region, as WUS directly promotes *JAZ3* expression in the SAM. Of note, JA-responsive *JAZ* genes are directly activated by MYC transcription factors and newly synthesized JAZ proteins in turn repress MYC activity, forming a negative feedback loop that attenuates JA signaling and limits its detrimental impact on plant growth and fitness. Since both MYC3 and WUS promote *JAZ3* expression, the JA responses are likely eliminated more rapidly within the *WUS*-expression domain compared to surrounding regions.

The interaction between WUS and JAZ3 further illustrates the multifaceted role of WUS in JA signaling regulation. WUS on the one hand directly activates the expression of *JAZ3* resulting in largely overlapping expression domains in the SAM and on the other hand WUS competes with COI1 for binding to JAZ3, thereby potentially preventing JAZ3 degradation. Stabilized JAZ3 represses JA signaling activation, thereby prioritizing growth-related processes under non-stressed conditions. However, our results show that wounding significantly increases JA and JA-Ile levels in both WT and *clv3-10* mutant, suggesting that WUS-mediated repression of JA signaling via JAZ3 is compromised under stress. It is therefore reasonable to propose that elevated JA concentrations favor the formation of JAZ3-COI1 complexes over JAZ3-WUS interaction, thereby shifting the regulatory balance from growth toward defense in response to systemic wounding. In line with this model, our yeast three-hybrid assays demonstrated that increasing JA-lle levels enhanced JAZ3-COI1 interaction even in the presence of WUS. These results indicate that JA-Ile concentration is an important signaling input of the WUS-JAZ3-COI1 interaction module, which orchestrates the growth-defense balance in the SAM under wounding stress. Intriguingly, previous studies have shown that SQUAMOSA PROMOTER BINDING PROTEIN-LIKE 9 (SPL9) similarly interferes with COI1-mediated JAZ3 degradation by interacting with JAZ3, thereby attenuating JA response in Arabidopsis leaves^48^. WUS-mediated JA attenuation in the SAM and SPL9-mediated regulation in the leaf both highlight a common strategy in which JAZ proteins serve as key regulatory nodes integrating JA signaling with developmental cues across different tissues.

In our study, we further reveal the potential to uncouple the growth-defense trade-off by genetically modulating the stem cell regulatory system. *Clv3-10* mutants exhibit both enhanced reproductive capacity and increased tolerance to stem growth inhibition caused by systemic wounding, suggesting that enhancing stem cell activity in the SAM can simultaneously promote growth and improve resilience to systemic wounding stress. Similarly, transgenic lines with elevated JAZ3 levels in the organizing center (*pWUS:JAZ3*) are more resistant to wounding-induced stem growth inhibition without compromising the overall inflorescence growth or developmental integrity. These findings suggest that the meristem represents a promising target for improving both plant productivity and stress tolerance in crop breeding. Future studies will be needed to further elucidate the mechanisms by which the WUS-JAZ3 module controls the growth–defense balance under various stress conditions.

## Methods

### Plant materials and growth conditions

All *Arabidopsis thaliana* plants used in this study were in the Columbia (Col-0) background, except for *wus-7*, which was in the Landsberg *erecta* (L*er*) background. The *clv3-10*, *wus-7, coi1-21*, *aos1-1* (SALK_017756), *jaz3/5/10* (SALK_139337, SALK_053775, SAIL_1156_C06), *pCOI1:COI1-VENUS, pUBQ10:mCherry-GR-WUS, pUBQ10:mCherry-GR,* and the triple reporter line (*TR7*) have been reported previously^34–38, 40, 43^. To generate *jaz3-cr* mutant, *pWUS:JAZ3*, and *pJAZ3:GFP-NLS* transgenic plants, corresponding constructs were assembled using GreenGate cloning system and transformed into plants via floral dip method. Arabidopsis plants were grown at 22 °C under long-day conditions (16 h light/8 h dark). *N. benthamiana* plants were grown at 25 °C under the same photoperiod. For seed germination, seeds were sterilized in 70% ethanol for 20 min, rinsed with 100% ethanol, and sowed on half-strength Murashige and Skoog (1/2 MS) medium containing 0.8% phyto agar. After stratification at 4 °C in the dark for two days, plates were transferred to growth chambers. Seedlings were transplanted to soil one week post-germination.

### Cloning

For yeast two-hybrid assays, the coding DNA sequences of *JAZ3* and *WUS* were amplified and cloned into pGADT7 and pGBKT7 vectors (Clontech), respectively. *JAZ3* was inserted using the *Eco*RI and *Xho*I restriction sites, and *WUS* using *Eco*RI and *Bam*HI. To generate pBridge_COI1_Em for yeast three-hybrid assays, the coding sequence of *COI1* was amplified and cloned into pBridge vector (Clontech) using *Sma*I and *Sal*I sites. For pBridge_COI1_WUS and pBridge_COI1_HEC1, the coding sequences of *WUS* and *HEC1* were amplified and cloned into pBridge_COI1_Em vectors using *Not*I and *Bgl*II sites. CRISPR/Cas9 constructs for gene knockout were generated as previously described^42^. Two gRNAs targeting the gene of interest were designed using the CHOP CHOP website ^(^http://chopchop.cbu.uib.no). To generate JAZ3_R305A/K306A_, site-directed mutagenesis was performed on the C module of *JAZ3* using the Quick-Change II XL Site-Directed Mutagenesis Kit (Agilent Technologies, 200521). Plasmids for other experiments were constructed using the GreenGate cloning system^20^. Details of modules and constructs are summarized in Table S1. Primer sequences used for cloning are listed in Table S2.

### Wounding and Chemical treatments

Wounding treatments were performed on rosette leaves as described in Fig. 1a. For quantification of inflorescence stem length, plants were wounded once a day for six consecutive days, the inflorescence stem length was measured on the seventh day. For imaging the SAMs of *TR7* plants, wounding was performed every day until the inflorescence stem reached a length of 10-12 cm. For gene expression analysis and quantification of OPDA, JA, and JA-Ile levels, six leaves per plants were wounded twice, and samples were collected three hours after treatments. For JA treatments, (+/-)-jasmonic acid (Sigma, J2500) was used. Unless otherwise specified, the working concentration of JA in all treatments was 50 μM, which was diluted from 200 mM JA stock using 100% ethanol as solvent. Treatments began when the main inflorescence stem was 0-0.5 cm in length. JA was applied by directly spraying Milli-Q water containing Silwet L-77 and JA onto the shoot apex. Mock treatments followed the same protocol but without JA. For SAM imaging and stem length measurement, JA was applied twice daily with measurements taken on the final day. For WUS induction, 10 μM DEX solution was sprayed onto the inflorescence apex. Mock treatments followed the same procedure but without DEX.

### Transient coexpression assay in *N. benthamiana*

*Agrobacterium tumefaciens* strain ASE harboring constructs of interest was cultured overnight in liquid medium at 28 °C. Bacterial cells were collected by centrifugation and resuspended in infiltration buffer (10 mM MgCl_2_, 10 mM MES, 0.15 mM acetosyringone). The optical density of suspensions was adjusted to OD _600_ = 0.5-0.7 prior to infiltration. The mixtures containing appropriate construct combinations were infiltrated into the abaxial side of leaves from four- to five-week-old *N. benthamiana* plants.

### AP-FRET

Leaf disks from *N. benthamiana* plants were harvested two days post-agroinfiltration. Fluorescence intensities (I) of donors (GFP-fused proteins) were recorded before and after photobleaching the acceptor fluorophore. FRET efficiency was calculated according to the following equation^49^: E(%) =100 x (I_post-bleach_-I_pre-bleach_)/I_post-bleach_.

### Confocal microscopy and image analysis

For SAM imaging, the inflorescence apex was dissected and inserted in 2% agarose medium. For DAPI staining, the dissected meristem was incubated in 1mg/ml DAPI for 1 min. Confocal imaging was performed using a Nikon A1 confocal microscope equipped with a CFI Apo LWD25× water immersion objective (Nikon Instruments). The following settings were applied: Ex: 488.4 nm, Em: 500-550 nm for GFP; Ex: 561.9 nm, Em: 570-620 nm for mCherry; Ex: 404.6 nm, Em: 425-475 nm for BFP and DAPI. Image analysis of *TR7* plants was processed as described^36^.

### RNA-sequencing and bioinformatics

DEX treatment on SAMs of *pUBQ10:mCherry-GR-WUS* plants was performed when the inflorescence stem was 3-5 cm in length. SAMs were finely dissected two hours post-treatment, and approximately 200 SAMs were pooled for each biological replicate. Total RNA extraction and RNA-sequencing were performed as previously described^37^. Raw sequencing reads were quality-checked with FastQC (v0.12.1), and adapter sequences along with low-quality bases were trimmed using Trimmomatic (v0.39). The filtered reads were aligned to the Arabidopsis reference genome using STAR (version 2.7.10b), and aligned reads were counted using HTSeq (v0.11.3). For further analyses, only genes with at least 10 counts in at least 3 samples were kept. Differential expression analysis was carried out on R Studio using DESeq2 (v1.42.1), with genes showcasing an adjusted p-value (p_.adj_) < 0.05 and |Log_2_FC| > 0.5 considered differentially expressed. GO enrichment analysis was performed using the web-based tool PANTHER (v19.0) with the Fisher’s Exact test and False Discovery Rate correction.

### Analysis of OPDA, JA, and JA-Ile concentrations in the shoot apex

20-25 mg frozen shoot apexes were homogenized in 750 µl MTBE and extracted for 30 min at 4 °C, followed by incubation for 15 min in an ultrasonic ice-batch. After centrifugation (10 min, 4 °C), the supernatant was transferred into a new reaction tube, and an equal volume of 0.1% HCl was added to the MTBE extract. The mixture was incubated for 30 min at 4 °C and centrifuged again. The upper phase was then collected and dried in a vacuum concentration system (SpeedVac, Eppendorf). Immediately before analysis, dried samples were reconstituted in 50 µl 50% ethanol by vortexing for 3 min. Chromatographic conditions and mass spectrometry parameters were carried out as previously described^50^.

### eChIP-qPCR

eChIP experiments were performed as previously described with minor modifications^20^. 5-day-old seedlings of *pUBQ10:mCherry-GR-WUS* and *pUBQ10:mCherry-GR* were treated with 10 μM DEX in 1/2 MS liquid medium for four hours. For each biological replicate, 0.1g seedlings was crosslinked with 1% formaldehyde for 10 min. Chromatin was sheared to an average size of 200-600 bp using a Bioruptor Pico (Diagenode; 30 s on/30 s off, 6 cycles, high frequency setting). The post-sonicated lysates were incubated with binding control magnetic agarose beads (ChromoTek, bmab-20) for two hours. A 1% aliquot of the lysate was saved as input. Immunoprecipitation was performed overnight using RFP-Trap magnetic beads (ChromoTek, rtma-20). DNA was purified using iPure kit v2 (Diagenode, C03010015), and qPCR data was analyzed via the percent input method.

### In situ hybridization

In situ hybridizations were performed as previously described^37^. Briefly, DNA templates for RNA probes synthesis were amplified from the respective C module vectors using primers flanked by T7 and SP6 promoter sequences. Probes were synthesized and labeled with Digoxegenin (DIG) (Roche, 11175025910) by in vitro transcription. For hybridization, inflorescence apexes were fixed in FAA solution (50% ethanol, 5% glacial acetic acid, 3.7% formaldehyde) and transferred into a Leica Asp200 for the automated vacuum and paraffin infiltration. The embedded samples were sectioned at a thickness of 8 μm and mounted onto glass slides. Hybridization was performed at 55 °C overnight. Following hybridization, slides were incubated with anti-DIG antibody (Roche, 11093274910) for 90 min. Signals were detected using NBT-BCIP (Roche, 11681451001) as the substrate.

### RNA extraction and RT-qPCR analysis

Total RNA was isolated from the inflorescence apex using the RNeasy Plant Mini kit (Qiagen, 74904). 1 μg of total RNA was reverse transcribed into cDNA using HiScript® III RT SuperMix (Vazyme, R323) followed the manufacturer’s instructions. qPCR analysis was conducted on a qTOWER^3^ Real-time thermal cycler (Analytik Jena) with Luna^®^ Universal qPCR master Mix (New England Biolabs, M3003L). *PROTEIN PHOSPHATASE 2A SUBUNIT A3* (*PP2A*) was used as the internal reference gene for normalization. All qPCR primer sequences are listed in Table S2, and relative gene expression levels were calculated from three technical replicates per sample using the comparative Ct (2^-ΔΔCt^) method.

### Yeast hybrid assays

All constructs were transformed into *Saccharomyces cerevisiae* strain AH109 using the EZ-Yeast Transformation Kit (MP Biomedicals, 112100200). Co-transformation of bait and prey vectors was performed according to the manufacturer’s instructions. Transformants were initially selected on DDO agar plates (Takara Bio, 630317). Three independent colonies were picked for each biological replicate and spotted onto quadruple dropout (QDO) agar plates (Takara Bio, 630323) supplemented with X-α-Gal (Roth, 107021-38-5). Plates were incubated at 28 °C for three days before imaging. For yeast three-hybrid assays, transformants grown on DDO plates were successively restreaked on the TDO plates three times. The refreshed transformants were then inoculated in TDO liquid medium supplemented with 40 μM Coronatine (Sigma, C8115) and incubated at 28 °C for three days. The optical density at 660 nm (OD_660_) was recorded, and β-galactosidase activity was quantified using the Yeast Beta-Galactosidase Assay Kit (Thermo Scientific, 75768) following the manufacturer’s microplate assay protocol (stopped method). For the JA-lIe-dose dependent assay, refreshed transformants were spotted on QDO and QuDO plates containing different concentrations of JA-Ile (Cayman Chemical, Cay10740-10).

### TurboID-based proximity labeling

50 μM biotin was infiltrated into *N. benthamiana* leaves two days after agroinfiltration. Tissues were harvested 30 min post-biotin treatment and immediately snap-frozen in liquid nitrogen. For protein extraction, tissues were ground into fine powder, and total protein from 0.5-1 g frozen tissues were extracted. The lysate was incubated and subsequently centrifuged at 4 °C. The supernatant was incubated with GFP-Trap Magnetic Beads (ChromoTek, gtma-20) for 60 min at 4 °C. The precipitate was washed 5 times at 4 °C. For immunoblotting, the following primary and secondary antibodies were used: SA (Invitrogen, s911; 1:10000), mouse anti-GFP (Santa Cruz Biotechnology, sc-9996; 1:6000), mouse anti-mCherry (abcam, ab125096; 1:2000), rabbit anti-HA (abcam, ab9110; 1:6000), donkey anti-mouse IgG-HRP (Invitrogen, A16011; 1:10000), and mouse anti-rabbit IgG-HRP (Santa Cruz Biotechnology, sc-2357; 1:10000).

### Statistics and reproducibility

All statistical analysis and data visualizations were performed using R (4.2.1). The normality of data distribution for each dataset was assessed using the Shapiro-Wilk test. For normally distributed datasets, the two-tailed Students’ t-test, Welch’s t-test, one-way ANOVA, or two-way ANOVA with post-hoc Tukey’s honestly significant difference test for multiple comparisons was applied accordingly. For datasets that were not normally distributed, the Wilcoxon rank-sum test or Kruskal-Wallis test followed by Dunn’s test for post-hoc multiple comparisons was performed accordingly.

## Data availability

All biological materials are available from the corresponding author. RNA-seq have been deposited in the NCBI Gene Expression Omnibus (GEO) under the accession numbers GSE311282. Source data are provided with this paper. All other data supporting the findings of this study are available in this article and supplementary information.

## Acknowledgements

We thank Sheng-Kai Sun (Centre for Organismal Studies, Heidelberg University) for sharing the *aos1-1* seeds, Bingwei Yu (College of Horticulture, South China Agricultural University) for sharing the pBridge vector for yeast three-hybrid experiments. We acknowledge the technical and scientific support of the Metabolomics Core Technology Platform (MCTP) of Heidelberg University. MCTP is partially funded by the CellNetworks Core Technology Platform (CCTP) of Heidelberg University, which is supported in part by the Federal Ministry of Education and Research (BMBF) and the Ministry of Science Baden-Württemberg within the framework of the Excellence Strategy of the Federal and State Governments of Germany. This research was supported by Heidelberg University and the CRC 1101 consortium of the Deutsche Forschungsgemeinschaft (DFG).

## Author contributions

P.F. and J.U.L. designed the experiments and wrote the paper with input from all other authors. Y.M. performed the RNA-seq experiments. P.B. analyzed the RNA-seq data and performed bioinformatic analyses. C.W. developed and optimized the 3D image analysis pipeline. G.P. performed the mass spectrometry analysis. J.Z. and T.G. cloned constructs for yeast two-hybrid experiments. P.F. conducted all the other experiments.

## Figures and figure legends

**Extended Data Fig. 1.**
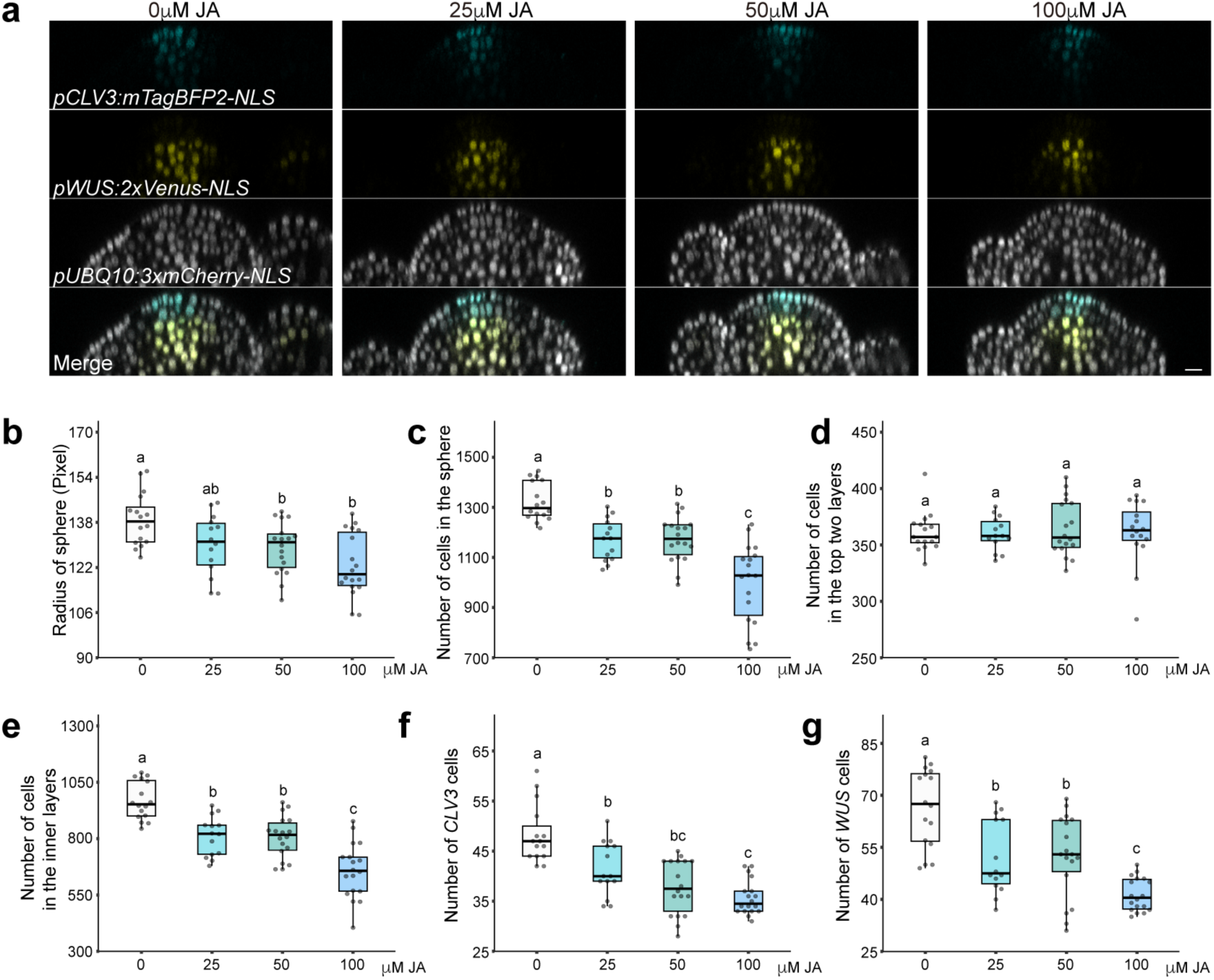
Dose-dependent effects of exogenous JA on SAM morphology. **a**, Representative central cross-sections of SAMs from the *TR7* triple reporter transgenic line after five days of treatment with different concentrations of JA. Scale bar, 10 μm. **b**-**g**, SAM morphology analysis of SAMs treated with different concentration of JA: **b**, Quantification of SAM size; **c**, Quantification of the number of cells in the SAM; **d**, Quantification of the number of cells in the top two layers of the SAM; **e**, Quantification of the number of cells in the inner layers of the SAM; **f**, Quantification of the number of *CLV3* cells. **g**, Quantification of the number of *WUS* cells. Each data point represents an independent measurement of the SAM from the main inflorescences stem. n = 14-18 per condition. Statistical significance was analyzed using one-way ANOVA followed by Tukey’s multiple comparisons test for **b**, **c**, **e** and **g**, and the Kruskal-Wallis test followed by Dunn’s multiple comparison test for **d** and **f** (*p* < 0.05).

**Extended Data Fig.2.**
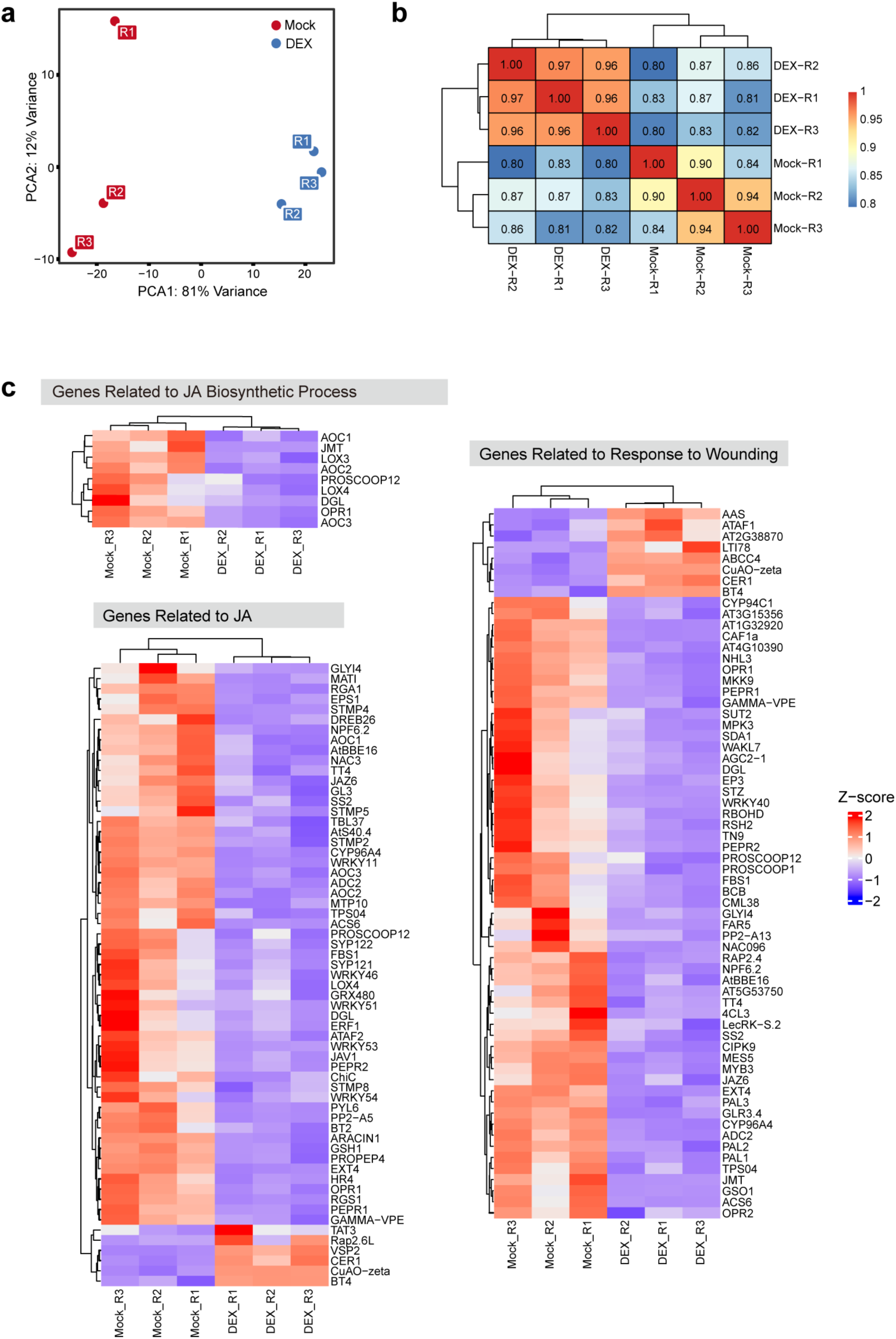
RNA-seq data from SAMs with ubiquitous induction of WUS activity. **a**, **b**, PCA plot (**a**) and Spearman Correlation Heatmap (**b**) of RNA-seq dataset from SAMs with inducible *WUS* overexpression. **c**, Heatmaps revealing expression of genes associated in JA or wounding pathways upon inducible WUS overexpression in the SAM. Genes showcasing an adjusted p-value (p._adj_) < 0.05 and |Log_2_FC|> 0.5 were analyzed and clustered based on enrichment of biologically relevant GO terms.

**Extended Data Fig.3.**
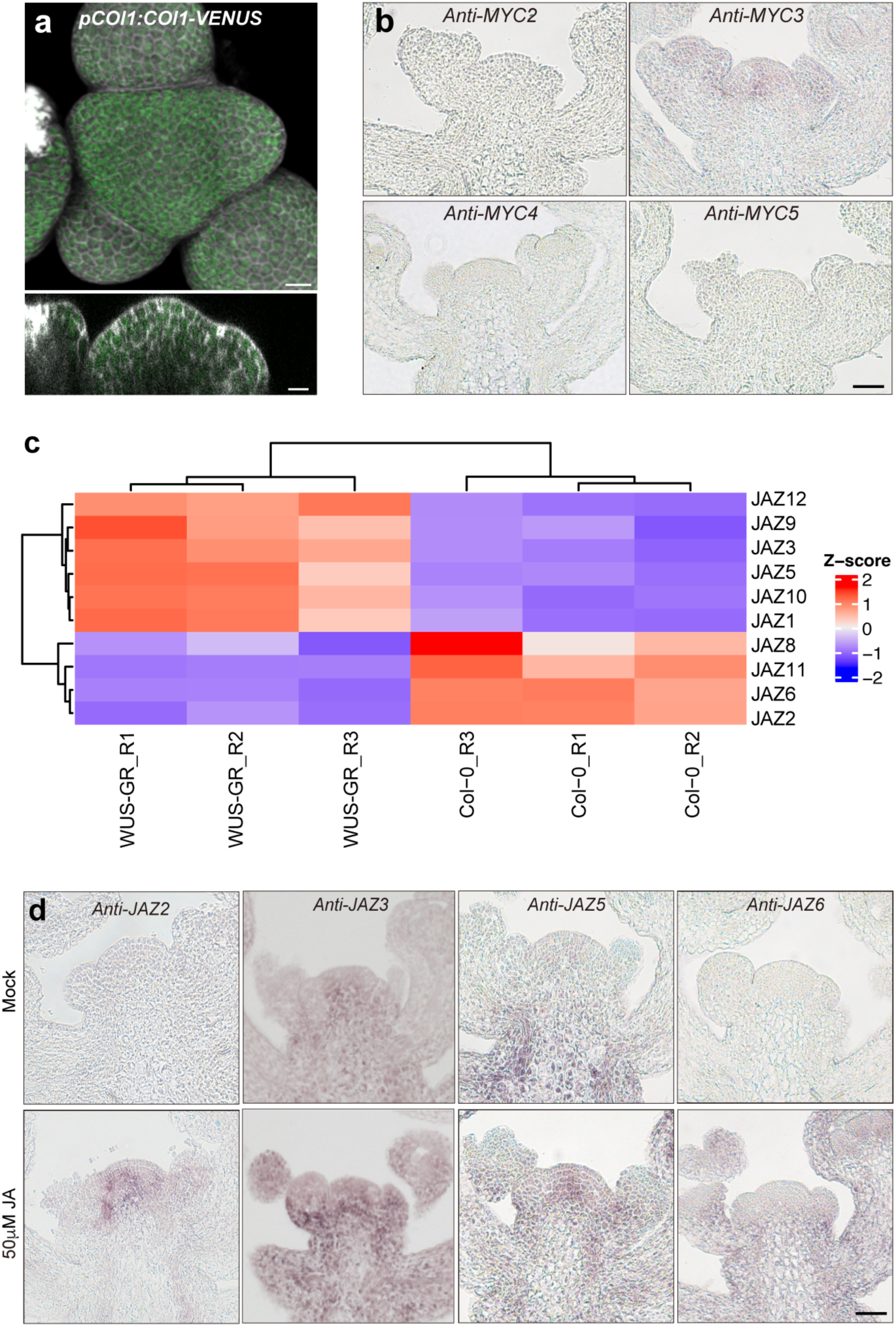
Expression pattern of representative components involved in JA signaling. **a**, Representative 3D volume view and central cross-section of the SAM in *pCOI1:COI1-VENUS* plants. Scale bar, 10 μm. The gray channel shows cell walls stained by DAPI. **b**, RNA accumulation patterns of *MYC2*, *MYC3*, *MYC4* and *MYC5* in the SAM detected by in situ hybridization. n = 4-10 per condition. **c**, Heatmaps revealing differential expression of *JAZ* genes in *pUBQ10:mCherry-GR-WUS* (WUS-GR) seedlings compared to WT(Col-0) upon DEX-induced WUS overexpression. Genes with an adjusted p-value (p_.adj_) < 0.05 and |Log_2_FC| > 0.5 were analyzed and presented in the heatmap. **d**, RNA accumulation patterns of *JAZ2*, *JAZ3*, *JAZ5*, and *JAZ6* in the SAM after two hours of mock or 50 μM JA treatment. n = 5-10. SAMs were treated and collected when the inflorescence stem length was 5-8 cm. Scale bar in **b** and **d**, 50 μm. Experiments were independently performed at least two times with similar results.

**Extended Data Fig.4.**
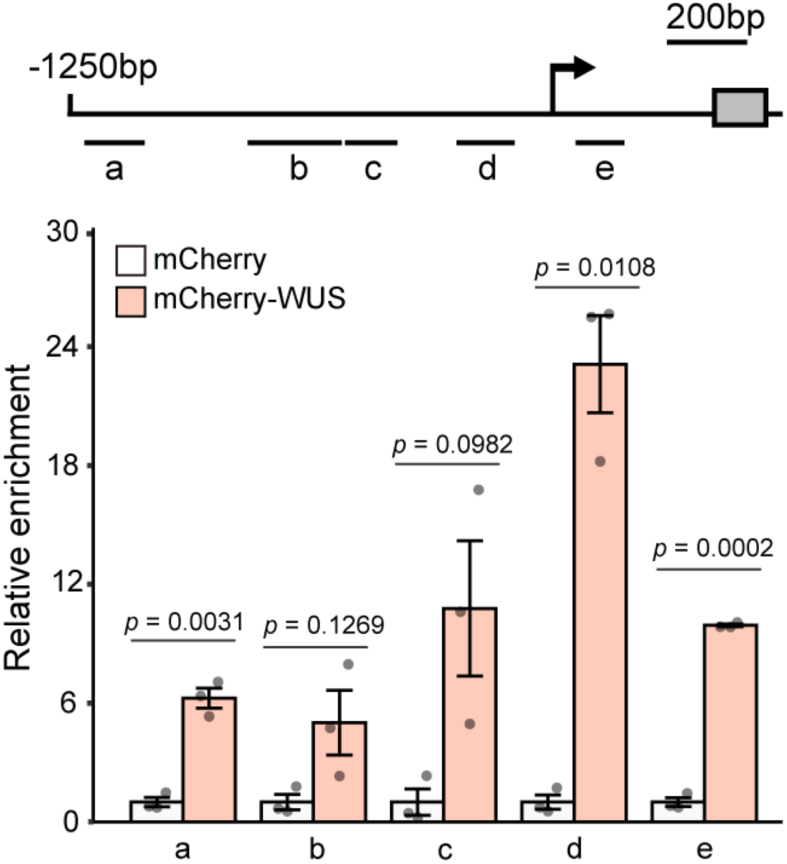
WUS binds to *JAZ3* promoter regions. Enrichments of *JAZ3* promoter and 5’-UTR fragments confirmed by eChIP-qPCR in *pUBQ10:mCherry-GR-linker-WUS* and *pUBQ10:mCherry-GR* seedlings with four-hour DEX treatment. In the upper panel, the black arrow marks the transcription start site and direction. The gray box indicates exon. Error bars indicate the mean ± s.d. of results from three biological replicates. *P* values were calculated using two-tailed Student’s t-test.

**Extended Data Fig.5.**
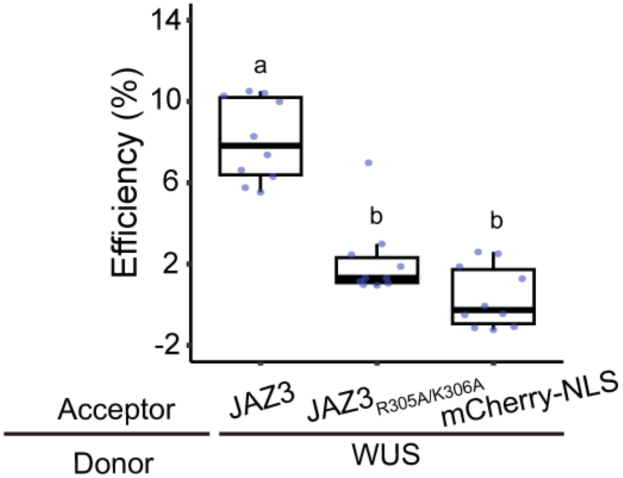
AP-FRET experiment reveals that two amino acids in Jas domain are essential for JAZ3 interacting with WUS. JAZ3_R305A/K306A_ is a JAZ3 mutant in which residues R305 and K306 are substituted with alanine. mCherry-JAZ3_R305A/K306A_ and WUS-GFP were used in this assay. Data points represent the FRET efficiencies measured from nuclei (n = 10). Statistical significance was analyzed using one-way ANOVA followed by Tukey’s multiple comparisons test (*p* < 0.05). Three biological replicates were performed with similar results.

**Extended Data Fig.6.**
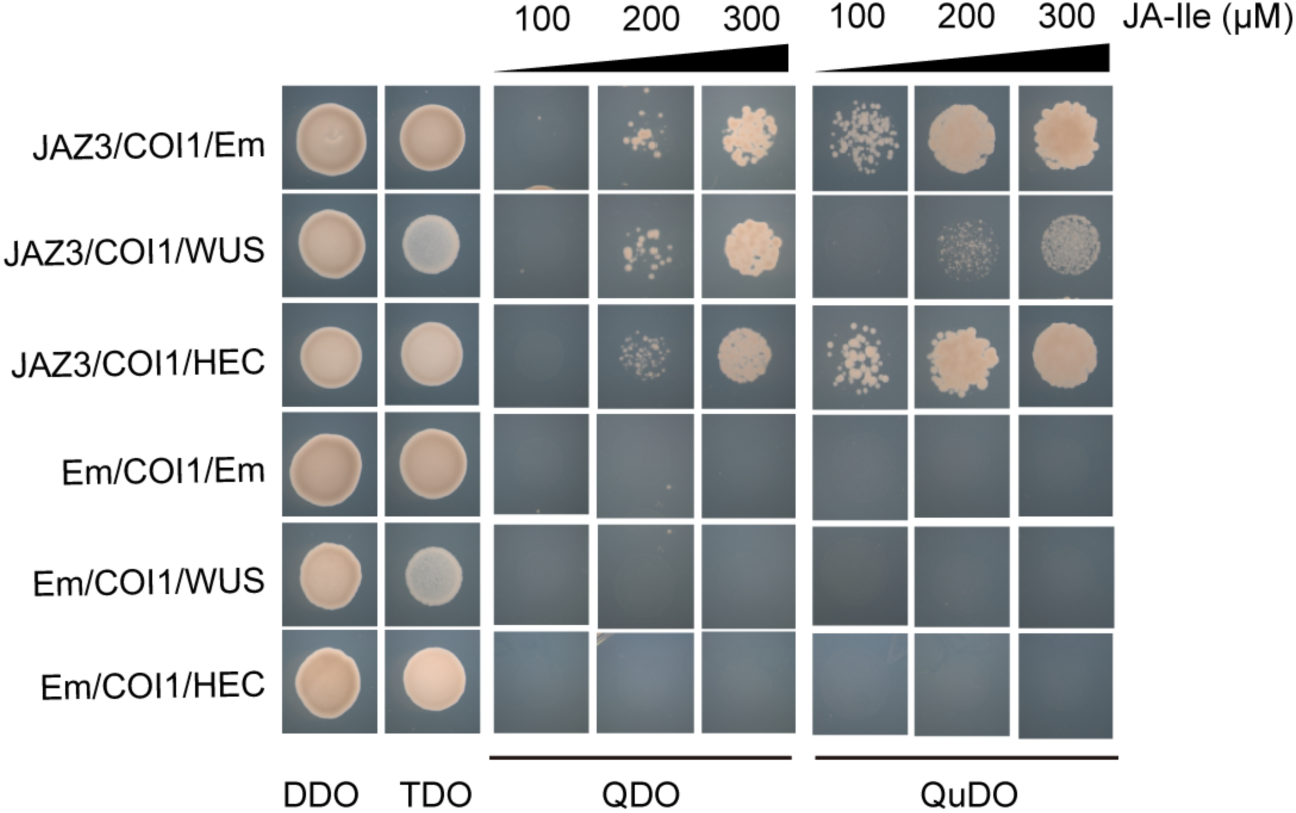
JA-dependent competition between WUS and COI1 for binding JAZ3 by Y3H assay. DDO (double dropout medium): SD/-Trp/-Leu. TDO (triple dropout medium): SD/-Trp/-Leu/-Met. QDO (quadruple dropout medium): SD/-Trp/-Leu/-His/-Ade. QuDO (Quintuple dropout medium): SD/-Trp/-Leu/-His/-Ade/-Met. Experiments were independently repeated twice with similar results.

**Extended Data Fig.7.**
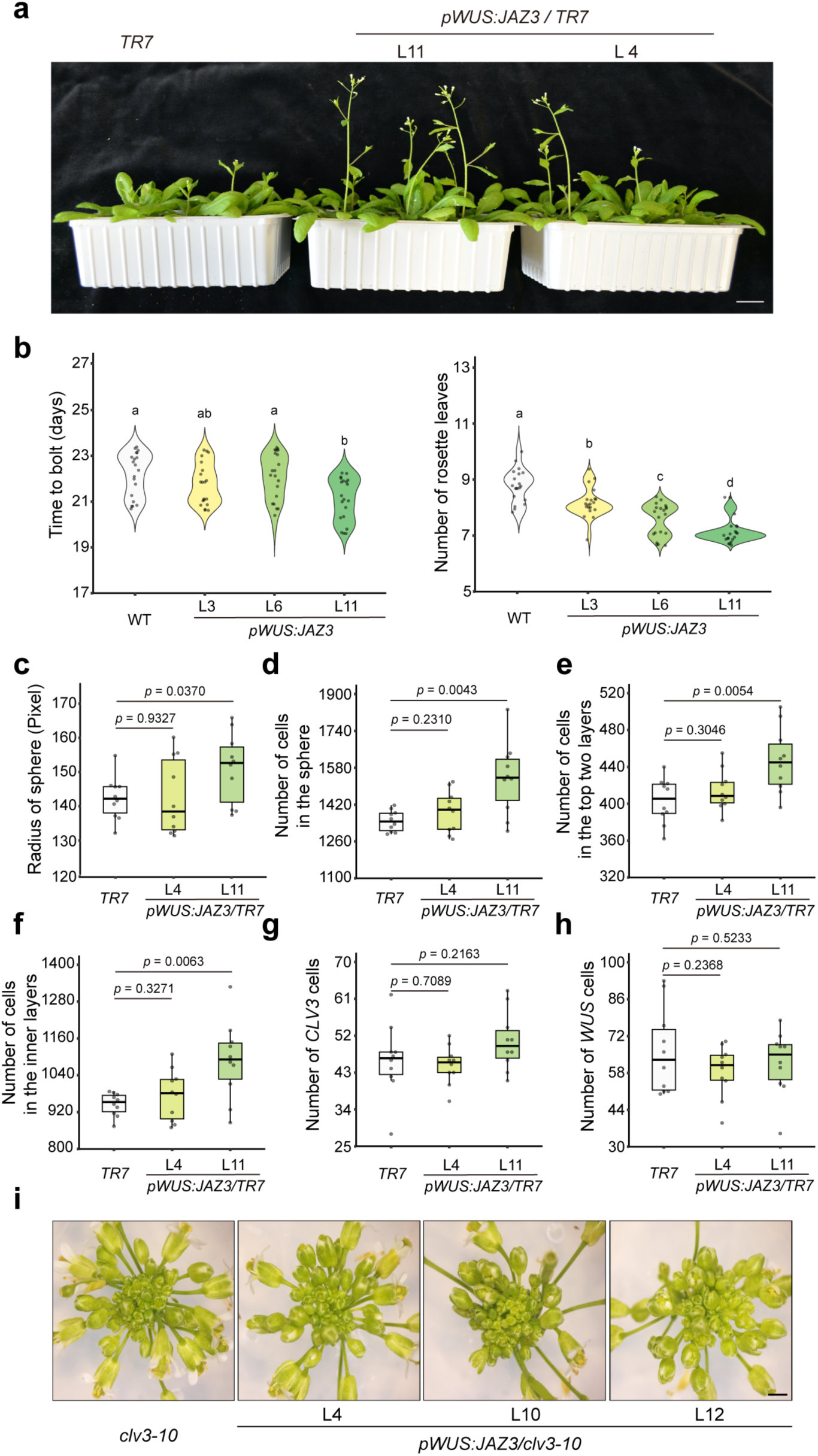
Developmental phenotypes of *pWUS:JAZ3* plants. **a**, Representative images of four-week-old *WUS:JAZ3* plants in *TR7* background. Scale bar, 2 cm. **b**, Quantification of time to bolt and number of rosette leaves at bolting in *pWUS:JAZ3* plants. Time to bolt (left) was recorded when the inflorescence stem length reached 0.5 cm, and the number of rosette leaves (right) was counted at the same time point. Data points represent results from one biological replicate (n = 20 for each condition). Statistical significances were analyzed using the Kruskal-Wallis test followed by Dunn’s multiple comparison test (P<0.05). Two biological replicates were performed with similar results. **c-h**, SAM morphology analysis of *pWUS:JAZ3* plants: **c**, Size of the defined SAM area; **d**, Total cell number within the defined SAM; **e**, Cell number of top two layers of the SAM; **f**, Cell number in the inner layers of the SAM; **g**, Cell number within *CLV3*-expressing domain. **h**, Cell number within the *WUS*-expressing domain. Each data point represents a measurement from the SAM of the main inflorescence meristem (n = 10 for each condition). The SAM was imaged when the inflorescence stem length was 10-12 cm. *P* values were calculated using two-tailed Student’s t-tests. **i**, the inflorescence tip of *clv3-10* and *pWUS:JAZ3/clv3-10* plants. Scale bar = 100 μm. The inflorescence tip was imaged when the inflorescence stem length of the plant was 8-10 cm.

**Extended Data Fig.8.**
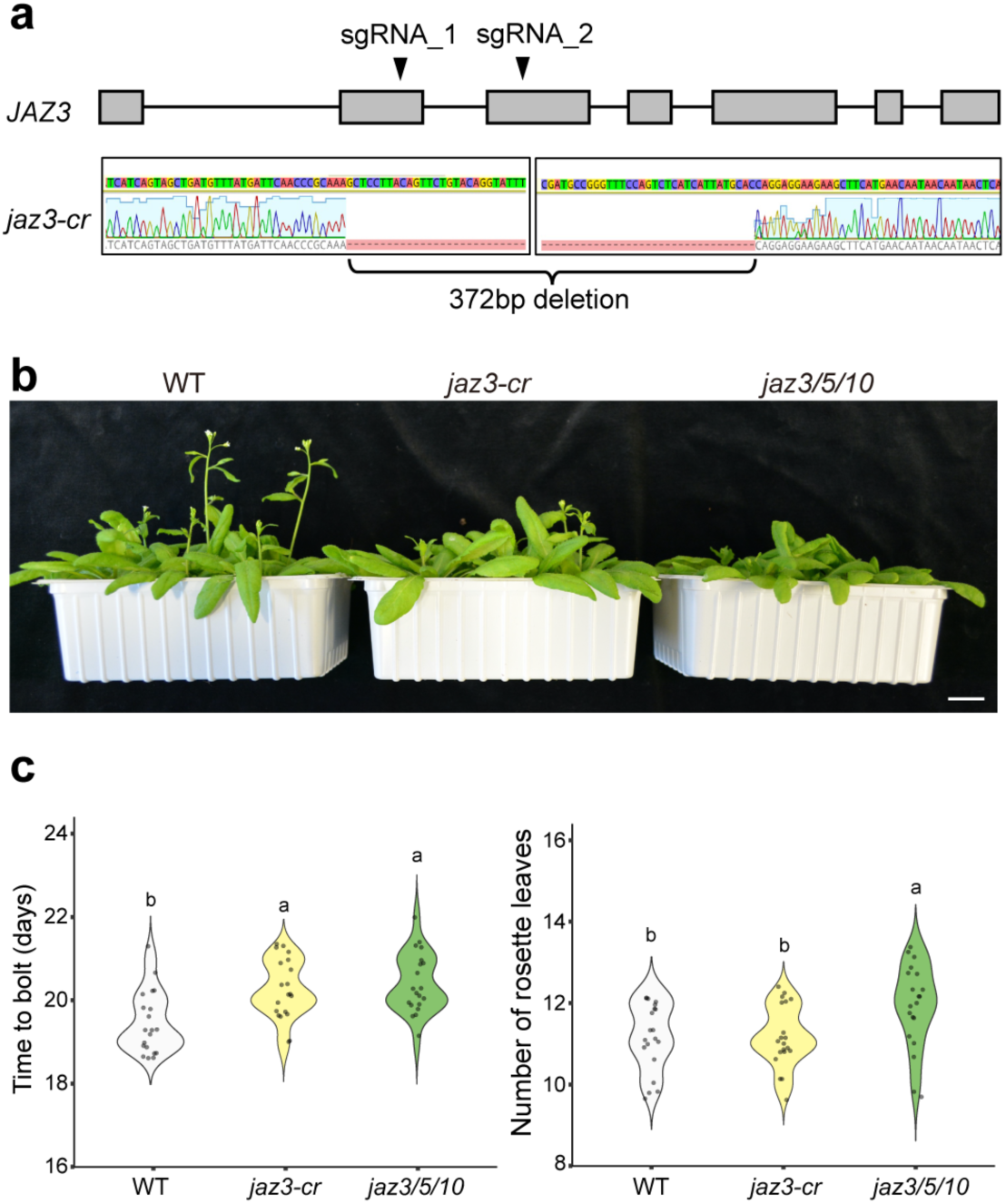
flowering phenotypes of *jaz3*-related mutants. **a**, Identification of *jaz3-cr* mutant generated by *CRISPR-Cas9*. The upper panel shows a schematic representation of the target sites of two sgRNAs within the *JAZ3* genomic region. Lower panel shows the resulting sequence deletion in the *JAZ3* locus. Grey boxes indicate exons, and black lines indicate introns. Arrowheads indicate the target sites of sgRNAs. **b**, Representative four-week-old WT, *jaz3-cr*, and *jaz3/5/10* mutant plants. Scale bar, 2 cm. **c**, Quantification of time to bolt and number of rosette leaves at bolting in *JAZ3*-related mutants. Time to bolt (left) was recorded when the inflorescence stem length reached 0.5 cm, and the number of leaves (right) was counted at the same time point. Data points represent results from one biological replicate (n = 20 for each condition). Statistical significances were analyzed using the Kruskal-Wallis test followed by Dunn’s multiple comparison test (P<0.05). Three biological replicates were performed with similar results.

## Supplementary tables

**Table. S1.**
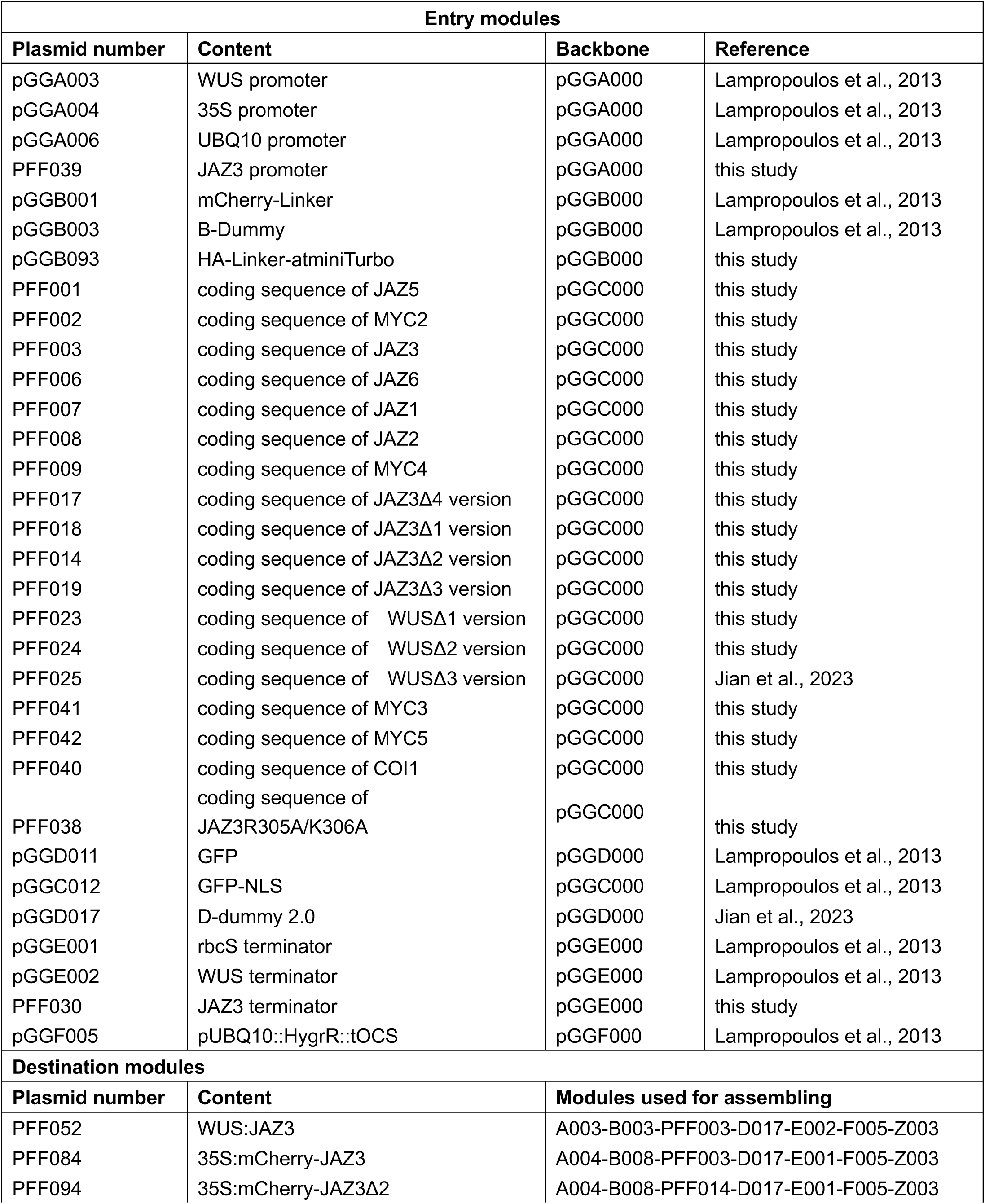
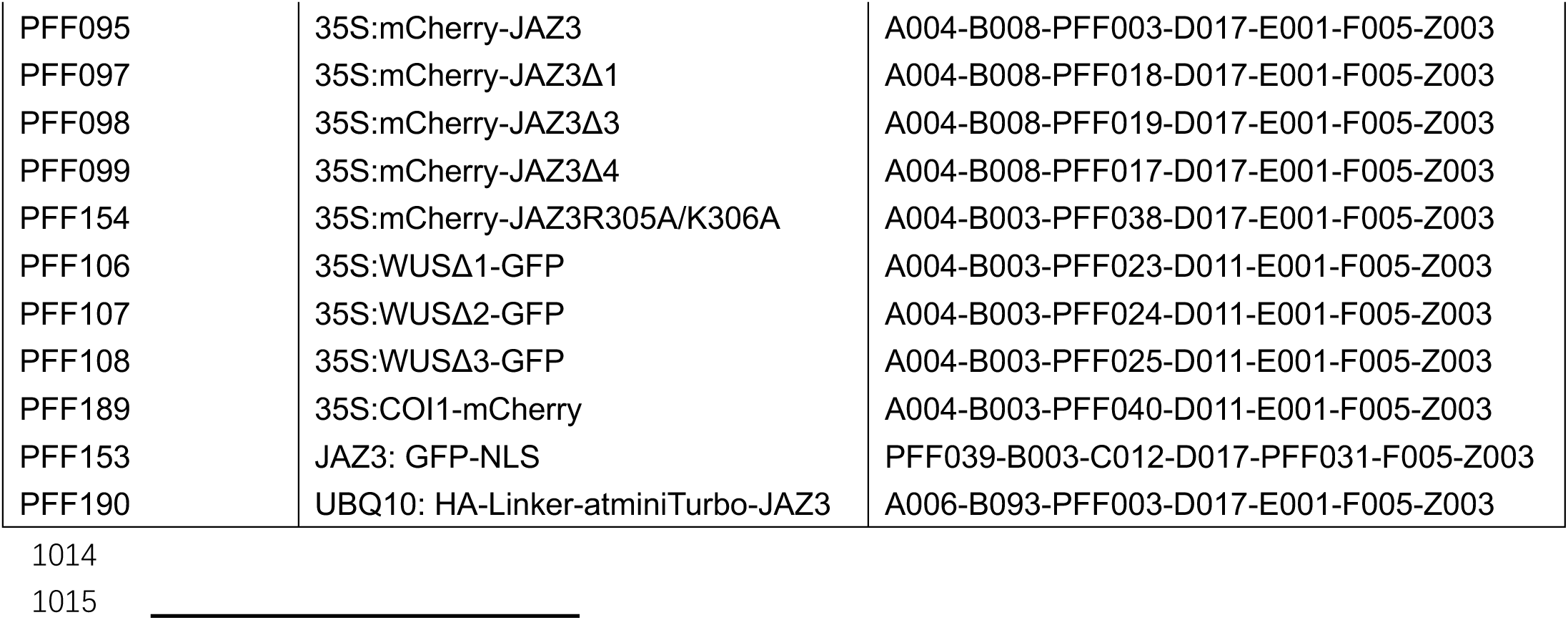
Plasmids used in this study.

**Table. S2.**
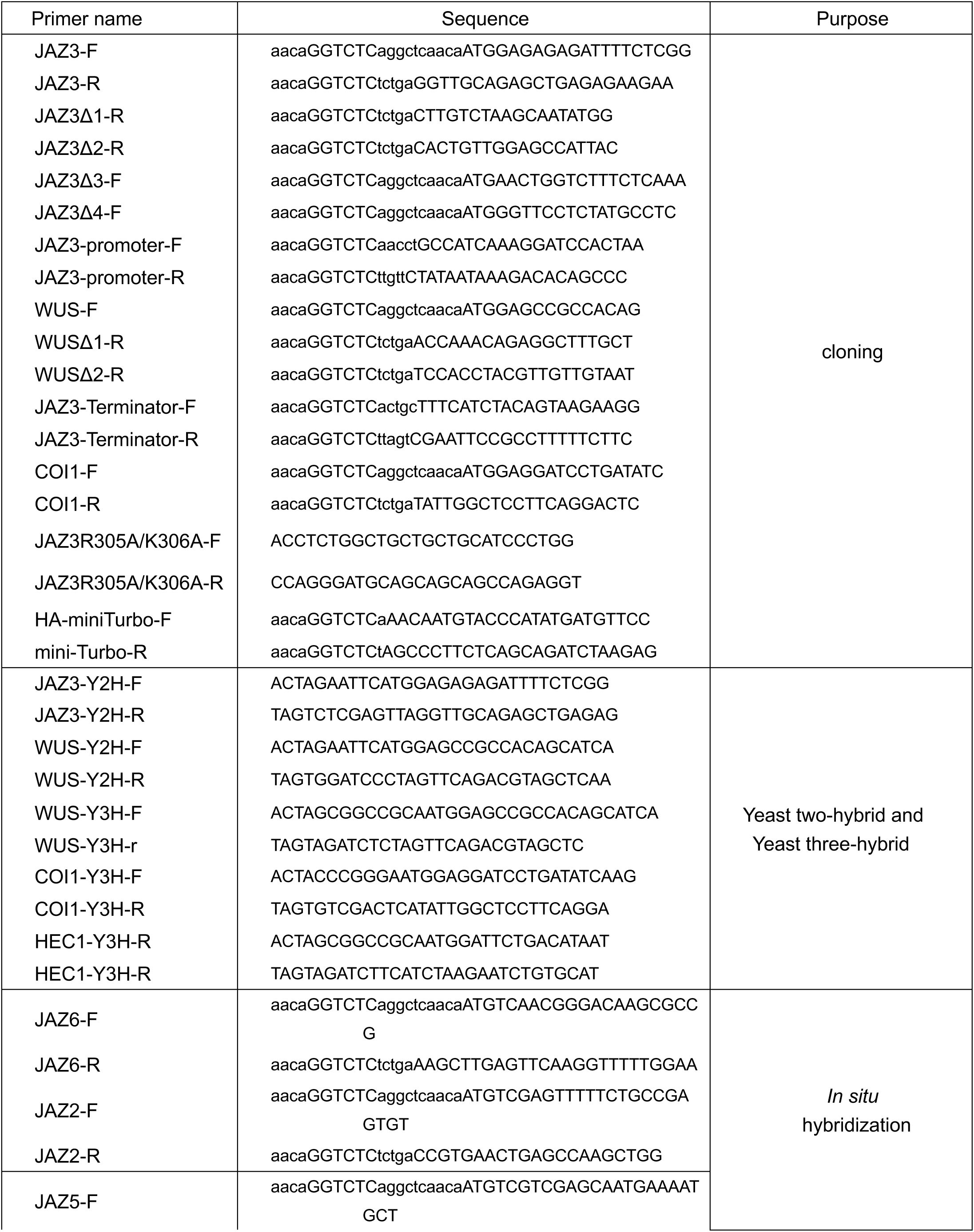
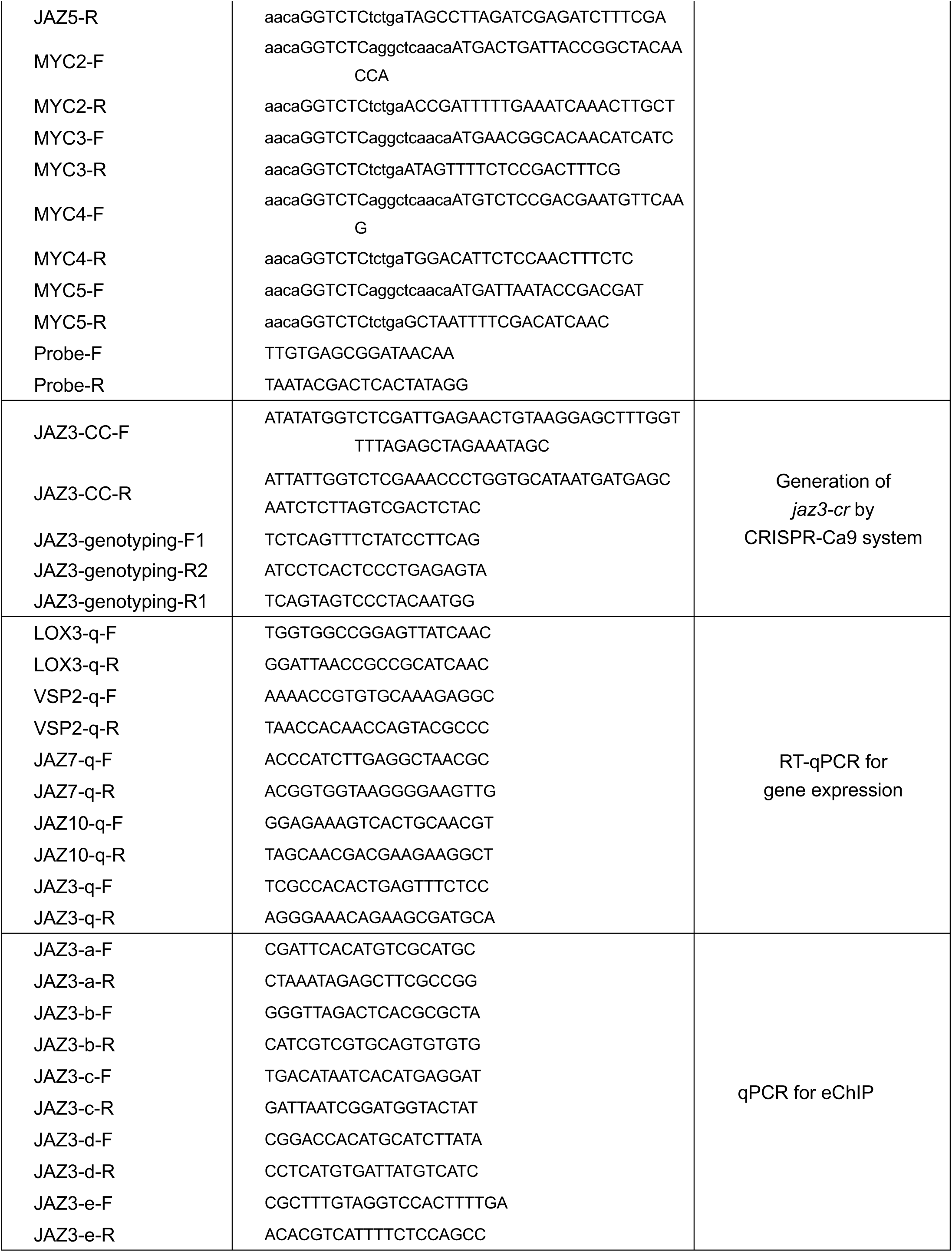
primers used in this study.

